## Supplementary figures and tables for "Complexes of vertebrate TMC1/2 and CIB2/3 proteins form hair-cell mechanotransduction cation channels"

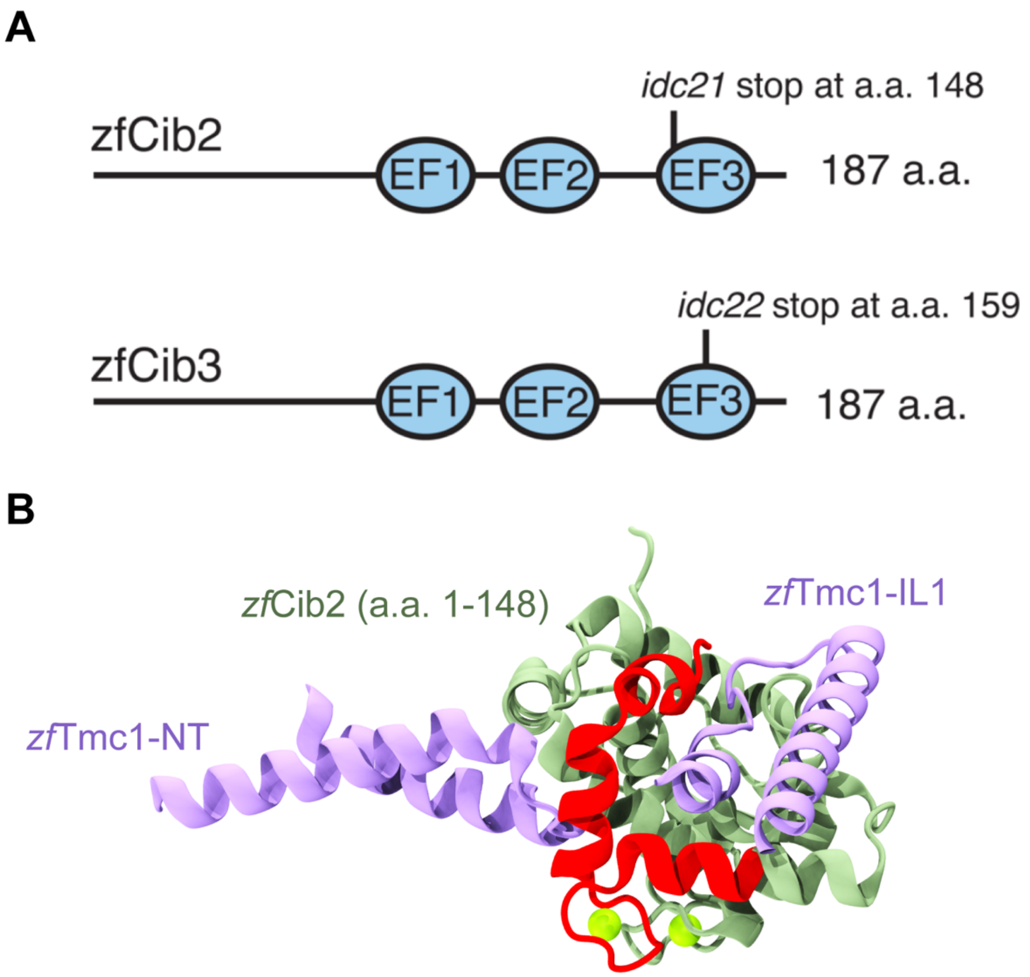


**Figure 5–figure supplement 1. Summary of zebrafish *cib2* and *cib3* alleles used in study.**
(A) Both the zebrafish Cib2 and Cib3 protein are 187 amino acids (a.a.) in length. We selected alleles of *cib2* and *cib3* that result in stop codons in the third EF-hand domain. In *cib2*$^{idc21}$ mutants a stop codon occurs at a.a. 148. In *cib3*$^{idc22}$ mutants a stop codon occurs at a.a. 159. (B) AF2 model showing zebrafish Tmc1-NT/IL1 (light purple), zebrafish Cib2 a.a. 1-148 (light green), and zebrafish Cib2 a.a. 149-187 (red). Position of $Ca^{2+}$ ions (green spheres) were obtained via superpostion of human CIB2 with zebrafish Cib2.

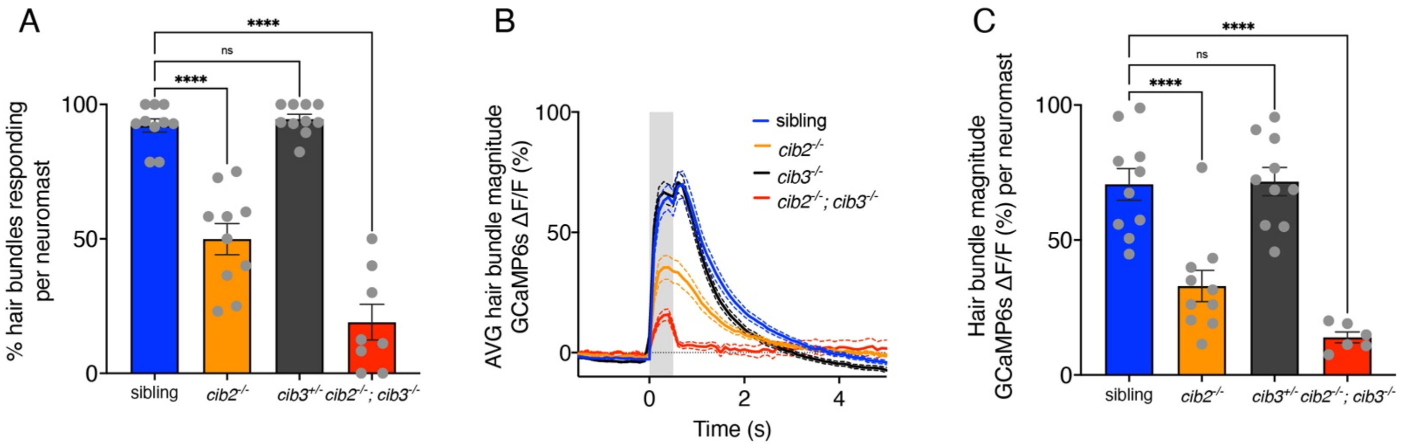


**Figure 5–figure supplement 2. Summary of mechanosensitive calcium responses in zebrafish.**
(**A**) Compared to controls, *cib3* mutants have a normal % of hair cells with mechanosensitive calcium responses per neuromast. In *cib2;cib3* mutants very few hair cells are mechanosensitive, and in *cib2* mutants there is a significantly reduced % of hair cells that are mechanosensitive per neuromast ($n$ = 10 neuromasts for sibling, *cib2*, and *cib3* mutants, and 8 neuromasts in *cib2;cib3* mutants). (**B**) Of the mechanosensitive hair bundles in (**A**), the average mechanosensitive calcium response is plotted for each genotype. (**C**) Of the mechanosensitive hair bundles in (**A**) the average response per neuromast is represented in a dot plot. Calcium imaging was performed at 5 or 6 dpf. A one-way ANOVA was used in **A** and **C**.

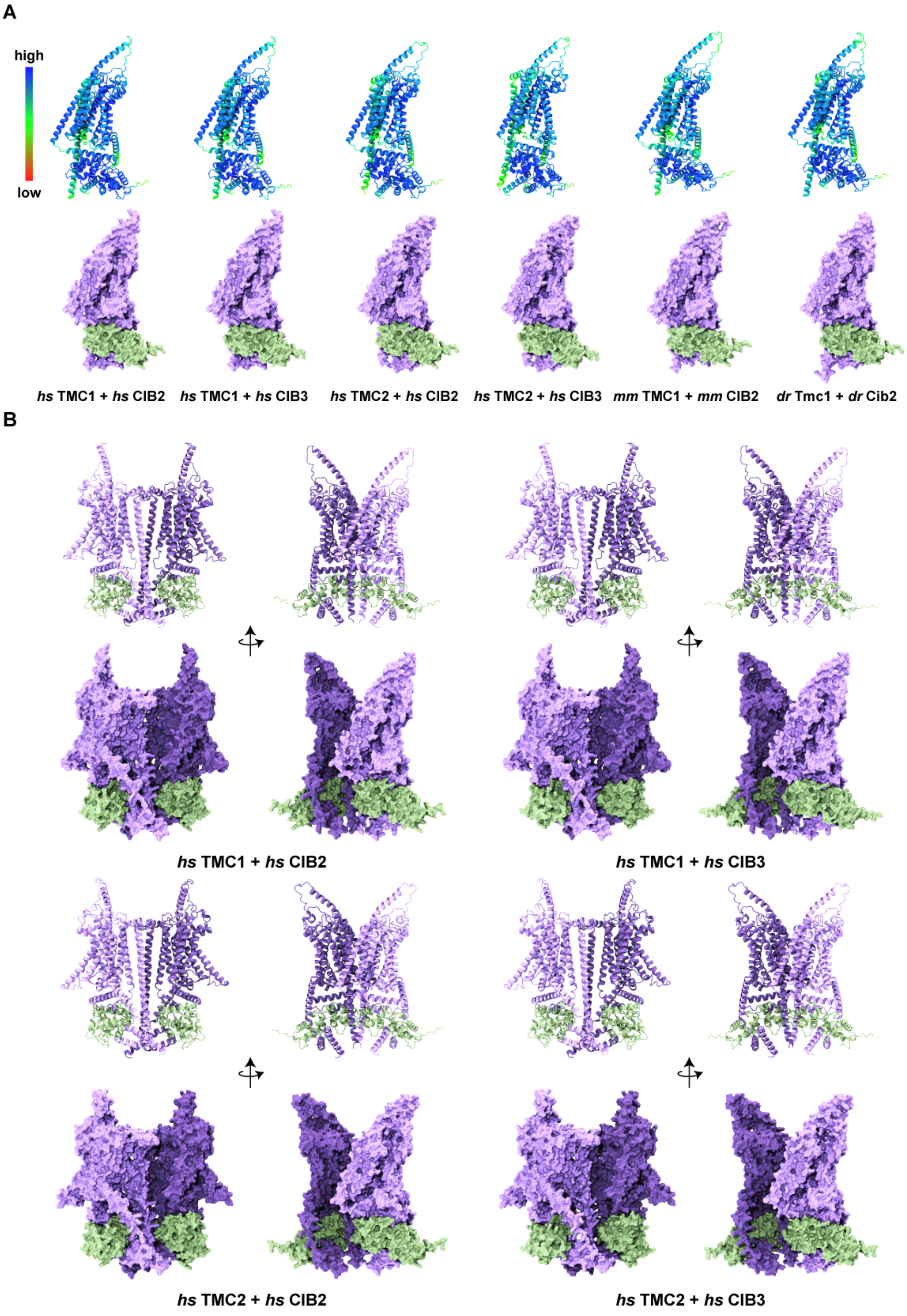


**Figure 7–figure supplement 1**. **Overview of TMC1/2 and CIB2/3 AF2 models.** (**A**) Human (*Homo sapiens*, *hs*), mouse (*Mus musculus*, *mm*), and zebrafish (*Danio rerio*, *dr*) models colored based on confidence scores are shown in the top row. Same models for TMC1/2 (light purple) and CIB2/3 (light sea green) are shown in surface representation on the bottom row. (**B**) Side views of homodimers of *hs* TMC1/2 (monomer A, light purple; monomer B, dark purple) in complex with *hs* CIB2/3 (light sea green) shown in ribbon and surface representation.

```
hs TMC1-1   -----------------------------------------------------
hs TMC1-2   -----------------------------------------------------
mm TMC1-1   -----------------------------------------------------
mm TMC1-2   -----------------------------------------------------
dr Tmc1-1   MLVKSELFMHATHHRQ----ALFSQAMVYCCDIIIIIIIIIIIITHTHGQPHA
dr Tmc1-2   ----METFNRKENHQEDLFLALFSQAMVYCCDIIIIIIIIIIIITHTHGQPHA
hs TMC2-1   -----------------------------------------------------
hs TMC2-2   -----------------------------------------------------
mm TMC2     -----------------------------------------------------
dr Tmc2a    -----------------------------------------------------
dr Tmc2b-1  -----------------------------------------------------
dr Tmc2b-2  -----------------------------------------------------
```

```
                             4    9
hs TMC1-1   -------------MSPKKVQIKVEEK---------------------------
hs TMC1-2   ------------MGKSKMIQIKVEEK---------------------------
mm TMC1-1   -----------------MLQIQVEEK---------------------------
mm TMC1-2   -----------------------------------------------------
dr Tmc1-1   RLLRRRMPRHKLIASESDVSIEVDEGKDKESCVYYVEVEENCERGKIKQASRD
dr Tmc1-2   RLLRRRMPRHKLIASESDVSIEVDEGKDKESCVYYVEVEENCERGKIKQASRD
hs TMC2-1   -------------MSHQVKGLKEEAR--------------GGVKGRVKSGS--
hs TMC2-2   ------MPRCHTLGCPVLSPAELTSS--RNHLGRLISHGRGHFPGPEAVGTRT
mm TMC2     -------------MSPQLKSLDEEGD---------------------------
dr Tmc2a    -----MPKKSDTTRKLEDVGIEIDGD--------------------VDSAEED
dr Tmc2b-1  -------------MCVLQLGWKWMQN---------------------------
dr Tmc2b-2  ----MPRRKSDIEITMDDVGMEMDAE---------------FDSGSDVDSKRR
```

```
                         14        24        34
hs TMC1-1   -------------EDETEESSSEEEEVEDKLPR--------------------
hs TMC1-2   -------------EDETEESSSEEEEVEDKLPR--------------------
mm TMC1-1   ---------------EEDTEESSSEEEEDKLPR--------------------
mm TMC1-2   -----------------------------------------------------
dr Tmc1-1   GKRRRERNGETRRKASEKRTNEGESKKAEKKHEKGHRTARKAGEKHGKRQRRK
dr Tmc1-2   GKRRRERNGETRRKASEKRTNEGESKKAEKKHEKGHRTARKAGEKHGKRQRRK
hs TMC2-1   ----PHTGDRLGRRSSSKRALKAEGTPGRRGAQRSQKERAGGSPSPGSPRR--
hs TMC2-2   VHFPAPAGDRLGRRSSSKRALKAEGTPGRRGAQRSQKERAGGSPSPGSPRR--
mm TMC2     ---------KSARRPTRKQTSRAACPQDGHRAQSSRKDPAKGSPRPGSSRK--
dr Tmc2a    KKSKGKGGKKAAGGKRGKASEDGEDEDEDDKPPKGRRAANKKKPAPVDEED--
dr Tmc2b-1  -----LIVEVMWTQKEEQKAGINQNLRREEGKPNQQTVWRRAKKRRMNRRA--
dr Tmc2b-2  AKGRNKPKPAARGRKAKPADSVEESEEEEDESPRKRRGRRKKQEDEDESEE--
```

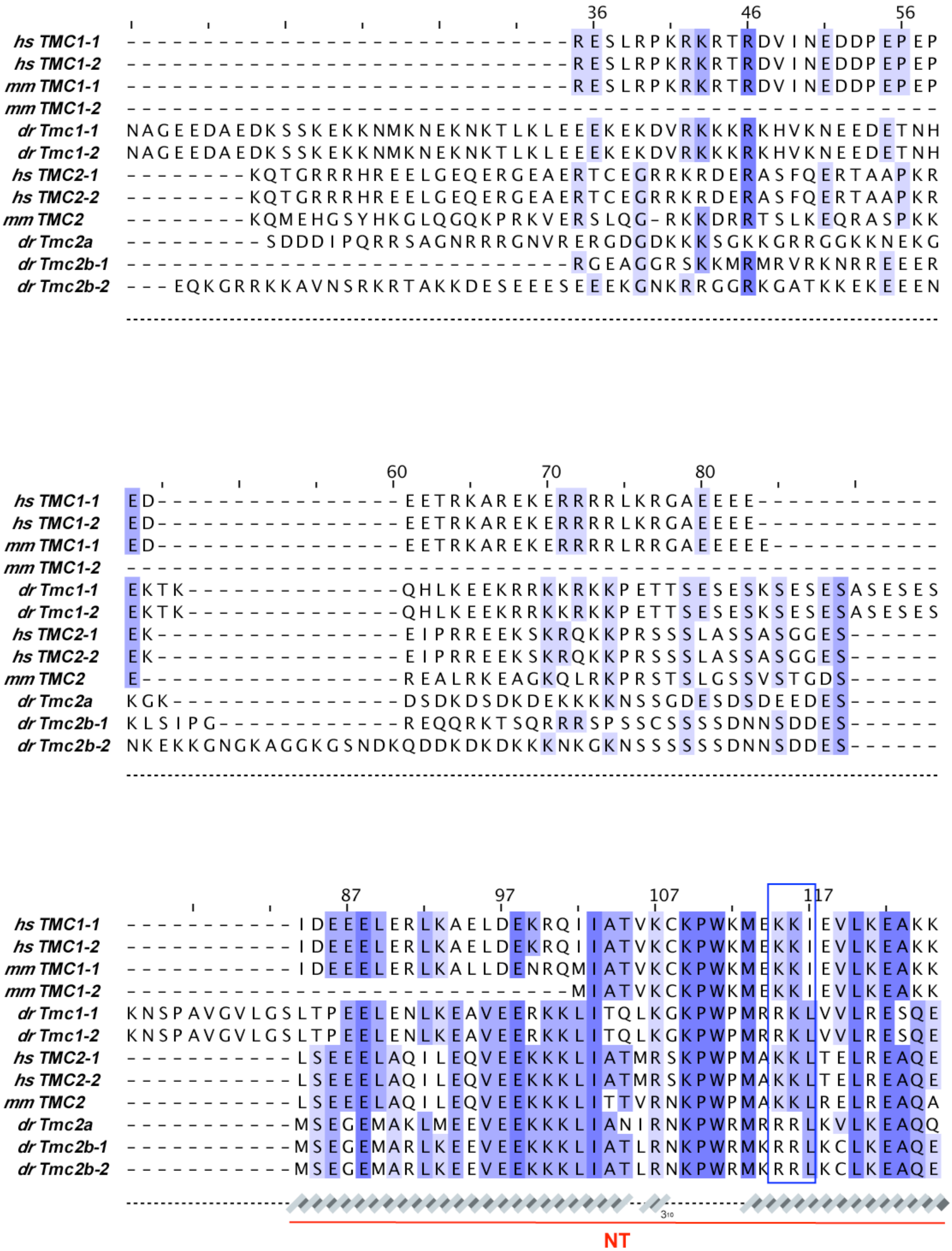
36
46
56
60
70
80
87
97
107
117
hs TMC1-1
hs TMC1-2
mm TMC1-1
mm TMC1-2
dr Tmc1-1
dr Tmc1-2
hs TMC2-1
hs TMC2-2
mm TMC2
dr Tmc2a
dr Tmc2b-1
dr Tmc2b-2
NT

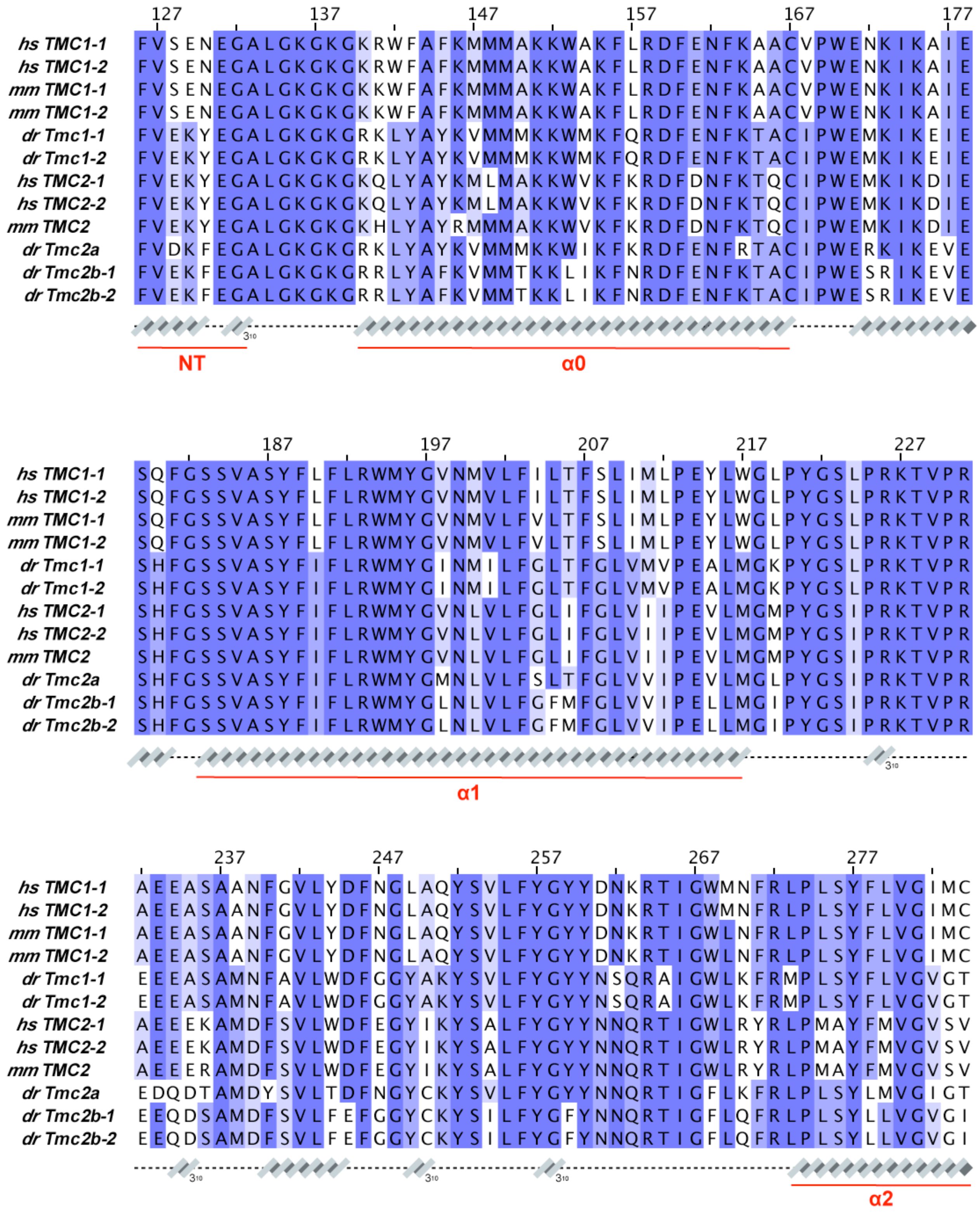
127 137 147 157 167 177
hs TMC1-1 FVSENEGALGKGKGKRWFAFKMMMAKKWAKFLRDFENFKAACVPWENKIKAIE
hs TMC1-2 FVSENEGALGKGKGKRWFAFKMMMAKKWAKFLRDFENFKAACVPWENKIKAIE
mm TMC1-1 FVSENEGALGKGKGKKWFAFKMMMAKKWAKFLRDFENFKAACVPWENKIKAIE
mm TMC1-2 FVSENEGALGKGKGKKWFAFKMMMAKKWAKFLRDFENFKAACVPWENKIKAIE
dr Tmc1-1 FVEKYEGALGKGKGRKLYAYKVMMMKKWMKFQRDFENFKTACIPWEMKIKEIE
dr Tmc1-2 FVEKYEGALGKGKGRKLYAYKVMMMKKWMKFQRDFENFKTACIPWEMKIKEIE
hs TMC2-1 FVEKYEGALGKGKGKQLYAYKMLMAKKWVKFKRDFDNFKTQCIPWEMKIKDIE
hs TMC2-2 FVEKYEGALGKGKGKQLYAYKMLMAKKWVKFKRDFDNFKTQCIPWEMKIKDIE
mm TMC2 FVEKYEGALGKGKGKHLYAYRMMMAKKWVKFKRDFDNFKTQCIPWEMKIKDIE
dr Tmc2a FVDKFEGALGKGKGRKLYAYKVMMMKKWIKFKRDFENFRTACIPWERKIKEVE
dr Tmc2b-1 FVEKFEGALGKGKGRRLYAFKVMMTKKLIKFNRDFENFKTACIPWESRIKEVE
dr Tmc2b-2 FVEKFEGALGKGKGRRLYAFKVMMTKKLIKFNRDFENFKTACIPWESRIKEVE
3₁₀
NT
α0
187 197 207 217 227
hs TMC1-1 SQFGSSVASYFLFLRWMYGVNMVLFILTFSLIMLPEYLWGLPYGSLPRKTVPR
hs TMC1-2 SQFGSSVASYFLFLRWMYGVNMVLFILTFSLIMLPEYLWGLPYGSLPRKTVPR
mm TMC1-1 SQFGSSVASYFLFLRWMYGVNMVLFVLTFSLIMLPEYLWGLPYGSLPRKTVPR
mm TMC1-2 SQFGSSVASYFLFLRWMYGVNMVLFVLTFSLIMLPEYLWGLPYGSLPRKTVPR
dr Tmc1-1 SHFGSSVASYFIFLRWMYGINMILFGLTFGLVMVPEALMGKPYGSLPRKTVPR
dr Tmc1-2 SHFGSSVASYFIFLRWMYGINMILFGLTFGLVMVPEALMGKPYGSLPRKTVPR
hs TMC2-1 SHFGSSVASYFIFLRWMYGVNLVLFGLIFGLVIIPEVLMGMPYGSIPRKTVPR
hs TMC2-2 SHFGSSVASYFIFLRWMYGVNLVLFGLIFGLVIIPEVLMGMPYGSIPRKTVPR
mm TMC2 SHFGSSVASYFIFLRWMYGVNLVLFGLIFGLVIIPEVLMGMPYGSIPRKTVPR
dr Tmc2a SHFGSSVASYFIFLRWMYGMNLVLFSLTFGLVVIPEVLMGLPYGSIPRKTVPR
dr Tmc2b-1 SHFGSSVASYFIFLRWMYGLNLVLFGFMFGLVVIPELLMGIPYGSIPRKTVPR
dr Tmc2b-2 SHFGSSVASYFIFLRWMYGLNLVLFGFMFGLVVIPELLMGIPYGSIPRKTVPR
3₁₀
α1
237 247 257 267 277
hs TMC1-1 AEEASAANFGVLYDFNGLAQYSVLFYGYYDNKRTIGWMNFRLPLSYFLVGIMC
hs TMC1-2 AEEASAANFGVLYDFNGLAQYSVLFYGYYDNKRTIGWMNFRLPLSYFLVGIMC
mm TMC1-1 AEEASAANFGVLYDFNGLAQYSVLFYGYYDNKRTIGWLNFRLPLSYFLVGIMC
mm TMC1-2 AEEASAANFGVLYDFNGLAQYSVLFYGYYDNKRTIGWLNFRLPLSYFLVGIMC
dr Tmc1-1 EEEASAMNFAVLWDFGGYAKYSVLFYGYYNSQRAIGWLKFRMPLSYFLVGVGT
dr Tmc1-2 EEEASAMNFAVLWDFGGYAKYSVLFYGYYNSQRAIGWLKFRMPLSYFLVGVGT
hs TMC2-1 AEEEKAMDFSVLWDFEGYIKYSALFYGYYNNQRTIGWLRYRLPMAYFMVGVSV
hs TMC2-2 AEEEKAMDFSVLWDFEGYIKYSALFYGYYNNQRTIGWLRYRLPMAYFMVGVSV
mm TMC2 AEEERAMDFSVLWDFEGYIKYSALFYGYYNNQRTIGWLRYRLPMAYFMVGVSV
dr Tmc2a EDQDTAMDYSVLTDFNGYCKYSVLFYGYYNNQRTIGFLKFRLPLSYLMVGIGT
dr Tmc2b-1 EEQDSAMDFSVLFEFGGYCKYSILFYGFYNNQRTIGFLQFRLPLSYLLVGVGI
dr Tmc2b-2 EEQDSAMDFSVLFEFGGYCKYSILFYGFYNNQRTIGFLQFRLPLSYLLVGVGI
3₁₀
3₁₀
3₁₀
α2

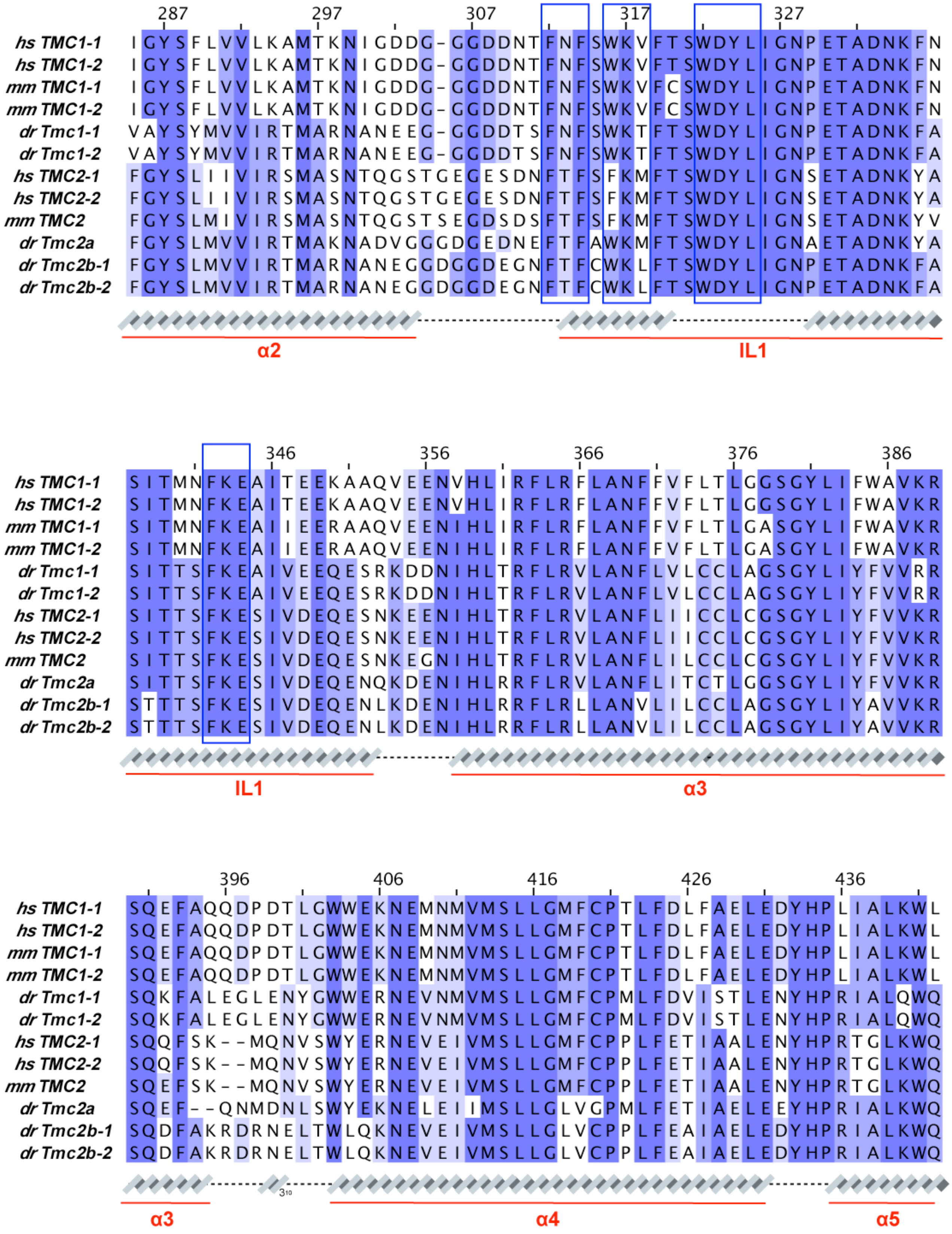
287 297 307 317 327
hs TMC1-1 IGYSFLVVLKAMTKNIGDDG-GGDDNTFNFSWKVFTSWDYLIGNPETADNKFN
hs TMC1-2 IGYSFLVVLKAMTKNIGDDG-GGDDNTFNFSWKVFTSWDYLIGNPETADNKFN
mm TMC1-1 IGYSFLVVLKAMTKNIGDDG-GGDDNTFNFSWKVFCSWDYLIGNPETADNKFN
mm TMC1-2 IGYSFLVVLKAMTKNIGDDG-GGDDNTFNFSWKVFCSWDYLIGNPETADNKFN
dr Tmc1-1 VAYSYMVVIRTMARNANEEG-GGDDTSFNFSWKTFTSWDYLIGNPETADNKFA
dr Tmc1-2 VAYSYMVVIRTMARNANEEG-GGDDTSFNFSWKTFTSWDYLIGNPETADNKFA
hs TMC2-1 FGYSLIIVIRSMASNTQGSTGEGESDNFTFSFKMFTSWDYLIGNSETADNKYA
hs TMC2-2 FGYSLIIVIRSMASNTQGSTGEGESDNFTFSFKMFTSWDYLIGNSETADNKYA
mm TMC2 FGYSLMIVIRSMASNTQGSTSEGDSDSFTFSFKMFTSWDYLIGNSETADNKYV
dr Tmc2a FGYSLMVVIRTMAKNADVGGGDGEDNEFTFAWKMFTSWDYLIGNAETADNKYA
dr Tmc2b-1 FGYSLMVVIRTMARNANEGGDGGDEGNFTFCWKLFTSWDYLIGNPETADNKFA
dr Tmc2b-2 FGYSLMVVIRTMARNANEGGDGGDEGNFTFCWKLFTSWDYLIGNPETADNKFA
α2
IL1
346 356 366 376 386
hs TMC1-1 SITMNFKEAITEEKAAQVEENVHLIRFLRFLANFFVFLTLGGSGYLIFWAVKR
hs TMC1-2 SITMNFKEAITEEKAAQVEENVHLIRFLRFLANFFVFLTLGGSGYLIFWAVKR
mm TMC1-1 SITMNFKEAIIEERAAQVEENIHLIRFLRFLANFFVFLTLGASGYLIFWAVKR
mm TMC1-2 SITMNFKEAIIEERAAQVEENIHLIRFLRFLANFFVFLTLGASGYLIFWAVKR
dr Tmc1-1 SITTSFKEAIVEEQESRKDDNIHLTRFLRVLANFLVLCCLAGSGYLIYFVVRR
dr Tmc1-2 SITTSFKEAIVEEQESRKDDNIHLTRFLRVLANFLVLCCLAGSGYLIYFVVRR
hs TMC2-1 SITTSFKESIVDEQESNKEENIHLTRFLRVLANFLIICCLCGSGYLIYFVVKR
hs TMC2-2 SITTSFKESIVDEQESNKEENIHLTRFLRVLANFLIICCLCGSGYLIYFVVKR
mm TMC2 SITTSFKESIVDEQESNKEGNIHLTRFLRVLANFLILCCLCGSGYLIYFVVKR
dr Tmc2a SITTSFKESIVDEQENQKDENIHLRRFLRVLANFLITCTLGGSGYLIYFVVKR
dr Tmc2b-1 STTTSFKESIVDEQENLKDENIHLRRFLRLLANVLILCCLAGSGYLIYAVVKR
dr Tmc2b-2 STTTSFKESIVDEQENLKDENIHLRRFLRLLANVLILCCLAGSGYLIYAVVKR
IL1
α3
396 406 416 426 436
hs TMC1-1 SQEFAQQDPDTLGWWEKNEMNMVMSLLGMFCPTLFDLFAELEDYHPLIALKWL
hs TMC1-2 SQEFAQQDPDTLGWWEKNEMNMVMSLLGMFCPTLFDLFAELEDYHPLIALKWL
mm TMC1-1 SQEFAQQDPDTLGWWEKNEMNMVMSLLGMFCPTLFDLFAELEDYHPLIALKWL
mm TMC1-2 SQEFAQQDPDTLGWWEKNEMNMVMSLLGMFCPTLFDLFAELEDYHPLIALKWL
dr Tmc1-1 SQKFALEGLENYGWWERNEVNMVMSLLGMFCPMLFDVISTLENYHPRIALQWQ
dr Tmc1-2 SQKFALEGLENYGWWERNEVNMVMSLLGMFCPMLFDVISTLENYHPRIALQWQ
hs TMC2-1 SQQFSK--MQNVSWYERNEVEIVMSLLGMFCPPLFETIAALENYHPRTGLKWQ
hs TMC2-2 SQQFSK--MQNVSWYERNEVEIVMSLLGMFCPPLFETIAALENYHPRTGLKWQ
mm TMC2 SQEFSK--MQNVSWYERNEVEIVMSLLGMFCPPLFETIAALENYHPRTGLKWQ
dr Tmc2a SQEF--QNMDNLSWYEKNELEIIMSLLGLVGPMLFETIAELEEYHPRIALKWQ
dr Tmc2b-1 SQDFAKRDRNELTWLQKNEVEIVMSLLGLVCPPLFEAIAELEDYHPRIALKWQ
dr Tmc2b-2 SQDFAKRDRNELTWLQKNEVEIVMSLLGLVCPPLFEAIAELEDYHPRIALKWQ
3₁₀
α3
α4
α5

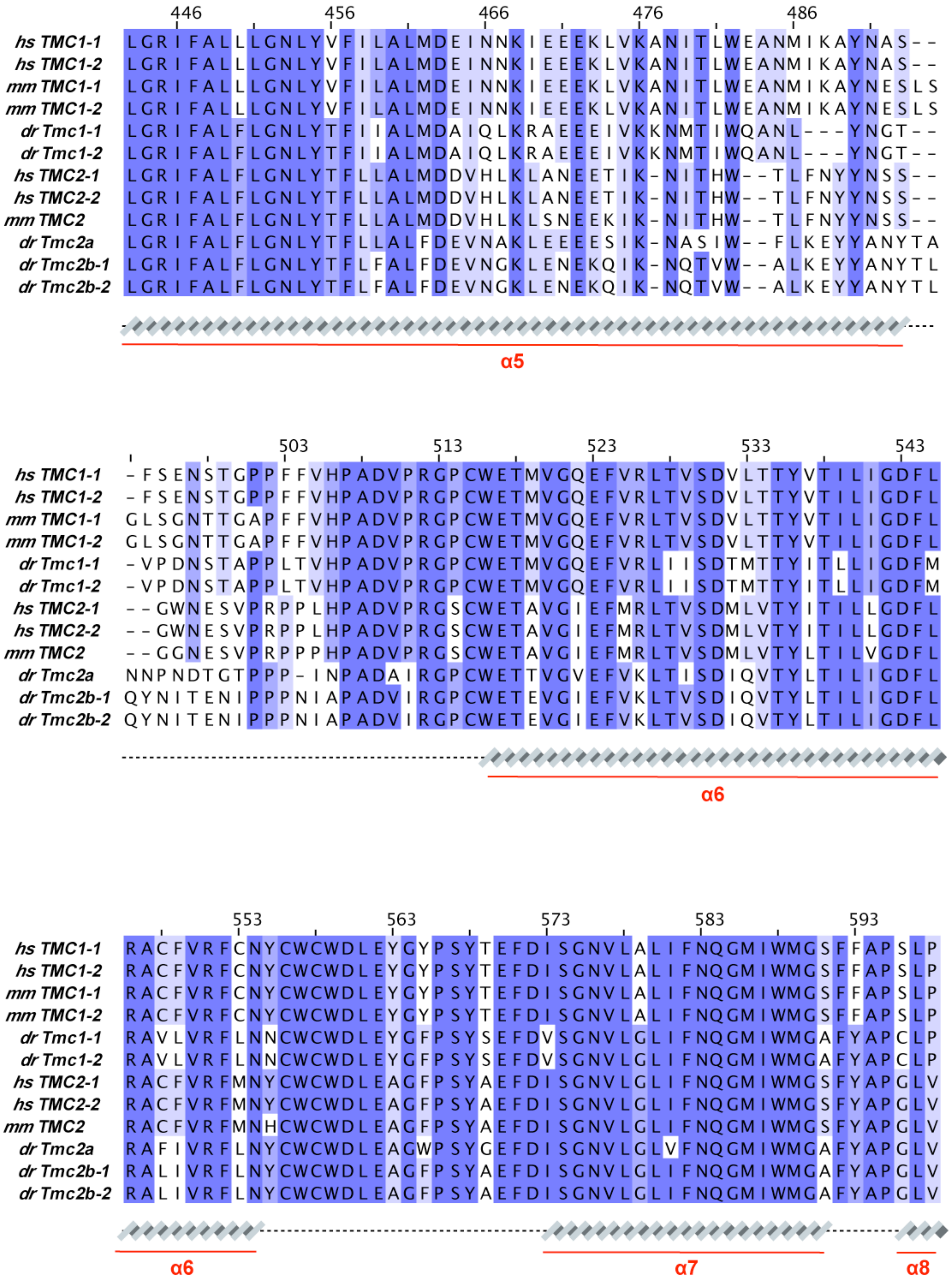
446 456 466 476 486
hs TMC1-1 LGRIFALLLGNLYVFILALMDEINNKIEEEKLVKANITLWEANMIKAYNAS--
hs TMC1-2 LGRIFALLLGNLYVFILALMDEINNKIEEEKLVKANITLWEANMIKAYNAS--
mm TMC1-1 LGRIFALLLGNLYVFILALMDEINNKIEEEKLVKANITLWEANMIKAYNESLS
mm TMC1-2 LGRIFALLLGNLYVFILALMDEINNKIEEEKLVKANITLWEANMIKAYNESLS
dr Tmc1-1 LGRIFALFLGNLYTFIIALMDAIQLKRAEEEIVKKNMTIWQANL---YNGT--
dr Tmc1-2 LGRIFALFLGNLYTFIIALMDAIQLKRAEEEIVKKNMTIWQANL---YNGT--
hs TMC2-1 LGRIFALFLGNLYTFLLALMDDVHLKLANEETIK-NITHW--TLFNYYNSS--
hs TMC2-2 LGRIFALFLGNLYTFLLALMDDVHLKLANEETIK-NITHW--TLFNYYNSS--
mm TMC2 LGRIFALFLGNLYTFLLALMDDVHLKLSNEEKIK-NITHW--TLFNYYNSS--
dr Tmc2a LGRIFALFLGNLYTFLLALFDEVNAKLEEEESIK-NASIW--FLKEYYANYTA
dr Tmc2b-1 LGRIFALFLGNLYTFLFALFDEVNGKLENEKQIK-NQTVW--ALKEYYANYTL
dr Tmc2b-2 LGRIFALFLGNLYTFLFALFDEVNGKLENEKQIK-NQTVW--ALKEYYANYTL
α5
503 513 523 533 543
hs TMC1-1 -FSENSTGPPFFVHPADVPRGPCWETMVGQEFVRLTVSDVLTTYVTILIGDFL
hs TMC1-2 -FSENSTGPPFFVHPADVPRGPCWETMVGQEFVRLTVSDVLTTYVTILIGDFL
mm TMC1-1 GLSGNTTGAPFFVHPADVPRGPCWETMVGQEFVRLTVSDVLTTYVTILIGDFL
mm TMC1-2 GLSGNTTGAPFFVHPADVPRGPCWETMVGQEFVRLTVSDVLTTYVTILIGDFL
dr Tmc1-1 -VPDNSTAPPLTVHPADVPRGPCWETMVGQEFVRLIISDTMTTYITLLIGDFM
dr Tmc1-2 -VPDNSTAPPLTVHPADVPRGPCWETMVGQEFVRLIISDTMTTYITLLIGDFM
hs TMC2-1 --GWNESVPRPPLHPADVPRGSCWETAVGIEFMRLTVSDMLVTYITILLGDFL
hs TMC2-2 --GWNESVPRPPLHPADVPRGSCWETAVGIEFMRLTVSDMLVTYITILLGDFL
mm TMC2 --GGNESVPRPPPHPADVPRGSCWETAVGIEFMRLTVSDMLVTYLTILVGDFL
dr Tmc2a NNPNDTGTPPP-INPADAIRGPCWETTVGVEFVKLTISDIQVTYLTILIGDFL
dr Tmc2b-1 QYNITENIPPPNIAPADVIRGPCWETEVGIEFVKLTVSDIQVTYLTILIGDFL
dr Tmc2b-2 QYNITENIPPPNIAPADVIRGPCWETEVGIEFVKLTVSDIQVTYLTILIGDFL
α6
553 563 573 583 593
hs TMC1-1 RACFVRFCNYCWCWDLEYGYPSYTEFDISGNVLALIFNQGMIWMGSFFAPSLP
hs TMC1-2 RACFVRFCNYCWCWDLEYGYPSYTEFDISGNVLALIFNQGMIWMGSFFAPSLP
mm TMC1-1 RACFVRFCNYCWCWDLEYGYPSYTEFDISGNVLALIFNQGMIWMGSFFAPSLP
mm TMC1-2 RACFVRFCNYCWCWDLEYGYPSYTEFDISGNVLALIFNQGMIWMGSFFAPSLP
dr Tmc1-1 RAVLVRFLNNCWCWDLEYGFPSYSEFDVSGNVLGLIFNQGMIWMGAFYAPCLP
dr Tmc1-2 RAVLVRFLNNCWCWDLEYGFPSYSEFDVSGNVLGLIFNQGMIWMGAFYAPCLP
hs TMC2-1 RACFVRFMNYCWCWDLEAGFPSYAEFDISGNVLGLIFNQGMIWMGSFYAPGLV
hs TMC2-2 RACFVRFMNYCWCWDLEAGFPSYAEFDISGNVLGLIFNQGMIWMGSFYAPGLV
mm TMC2 RACFVRFMNHCWCWDLEAGFPSYAEFDISGNVLGLIFNQGMIWMGSFYAPGLV
dr Tmc2a RAFIVRFLNYCWCWDLEAGWPSYGEFDISGNVLGLVFNQGMIWMGAFYAPGLV
dr Tmc2b-1 RALIVRFLNYCWCWDLEAGFPSYAEFDISGNVLGLIFNQGMIWMGAFYAPGLV
dr Tmc2b-2 RALIVRFLNYCWCWDLEAGFPSYAEFDISGNVLGLIFNQGMIWMGAFYAPGLV
α6
α7
α8

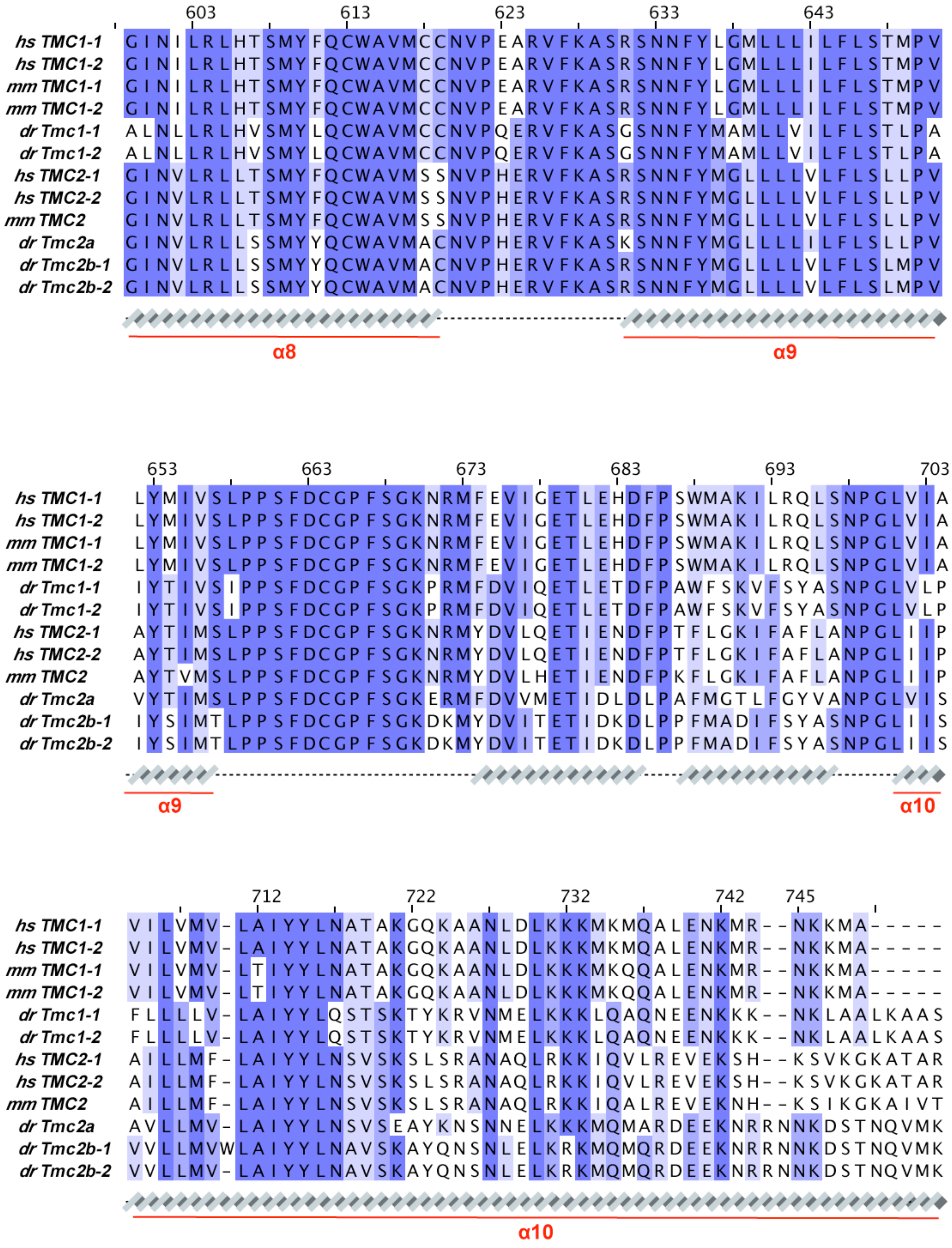

603 613 623 633 643
hs TMC1-1 GINILRLHTSMYFQCWAVMCCNVPEARVFKASRSNNFYLGMLLLILFLSTMPV
hs TMC1-2 GINILRLHTSMYFQCWAVMCCNVPEARVFKASRSNNFYLGMLLLILFLSTMPV
mm TMC1-1 GINILRLHTSMYFQCWAVMCCNVPEARVFKASRSNNFYLGMLLLILFLSTMPV
mm TMC1-2 GINILRLHTSMYFQCWAVMCCNVPEARVFKASRSNNFYLGMLLLILFLSTMPV
dr Tmc1-1 ALNLLRLHVSMYLQCWAVMCCNVPQERVFKASGSNNFYMAMLLVILFLSTLPA
dr Tmc1-2 ALNLLRLHVSMYLQCWAVMCCNVPQERVFKASGSNNFYMAMLLVILFLSTLPA
hs TMC2-1 GINVLRLLTSMYFQCWAVMSSNVPHERVFKASRSNNFYMGLLLLVLFLSLLPV
hs TMC2-2 GINVLRLLTSMYFQCWAVMSSNVPHERVFKASRSNNFYMGLLLLVLFLSLLPV
mm TMC2 GINVLRLLTSMYFQCWAVMSSNVPHERVFKASRSNNFYMGLLLLVLFLSLLPV
dr Tmc2a GINVLRLLSSMYYQCWAVMACNVPHERVFKASKSNNFYMGLLLLILFLSLLPV
dr Tmc2b-1 GINVLRLLSSMYYQCWAVMACNVPHERVFKASRSNNFYMGLLLLVLFLSLMPV
dr Tmc2b-2 GINVLRLLSSMYYQCWAVMACNVPHERVFKASRSNNFYMGLLLLVLFLSLMPV
α8
α9
653 663 673 683 693 703
hs TMC1-1 LYMIVSLPPSFDCGPFSGKNRMFEVIGETLEHDFPSWMAKILRQLSNPGLVIA
hs TMC1-2 LYMIVSLPPSFDCGPFSGKNRMFEVIGETLEHDFPSWMAKILRQLSNPGLVIA
mm TMC1-1 LYMIVSLPPSFDCGPFSGKNRMFEVIGETLEHDFPSWMAKILRQLSNPGLVIA
mm TMC1-2 LYMIVSLPPSFDCGPFSGKNRMFEVIGETLEHDFPSWMAKILRQLSNPGLVIA
dr Tmc1-1 IYTIVSIPPSFDCGPFSGKPRMFDVIQETLETDFPAWFSKVFSYASNPGLVLP
dr Tmc1-2 IYTIVSIPPSFDCGPFSGKPRMFDVIQETLETDFPAWFSKVFSYASNPGLVLP
hs TMC2-1 AYTIMSLPPSFDCGPFSGKNRMYDVLQETIENDFPTFLGKIFAFLANPGLIIP
hs TMC2-2 AYTIMSLPPSFDCGPFSGKNRMYDVLQETIENDFPTFLGKIFAFLANPGLIIP
mm TMC2 AYTVMSLPPSFDCGPFSGKNRMYDVLHETIENDFPKFLGKIFAFLANPGLIIP
dr Tmc2a VYTIMSLPPSFDCGPFSGKERMFDVVMETIDLDLPAFMGTLFGYVANPGLVIS
dr Tmc2b-1 IYSIMTLPPSFDCGPFSGKDKMYDVITETIDKDLPPFMADIFSYASNPGLIIS
dr Tmc2b-2 IYSIMTLPPSFDCGPFSGKDKMYDVITETIDKDLPPFMADIFSYASNPGLIIS
α9
α10
712 722 732 742 745
hs TMC1-1 VILVMV-LAIYYLNATAKGQKAANLDLKKKMKMQALENKMR--NKKMA-----
hs TMC1-2 VILVMV-LAIYYLNATAKGQKAANLDLKKKMKMQALENKMR--NKKMA-----
mm TMC1-1 VILVMV-LTIYYLNATAKGQKAANLDLKKKMKQQALENKMR--NKKMA-----
mm TMC1-2 VILVMV-LTIYYLNATAKGQKAANLDLKKKMKQQALENKMR--NKKMA-----
dr Tmc1-1 FLLLLV-LAIYYLQSTSKTYKRVNMELKKKLQAQNEENKKK--NKLAALKAAS
dr Tmc1-2 FLLLLV-LAIYYLQSTSKTYKRVNMELKKKLQAQNEENKKK--NKLAALKAAS
hs TMC2-1 AILLMF-LAIYYLNSVSKSLSRANAQLRKKIQVLREVEKSH--KSVKGKATAR
hs TMC2-2 AILLMF-LAIYYLNSVSKSLSRANAQLRKKIQVLREVEKSH--KSVKGKATAR
mm TMC2 AILLMF-LAIYYLNSVSKSLSRANAQLRKKIQALREVEKNH--KSIKGKAIVT
dr Tmc2a AVLLMV-LAIYYLNSVSEAYKNSNNELKKKMQMARDEEKNRRNNKDSTNQVMK
dr Tmc2b-1 VVLLMVWLAIYYLNAVSKAYQNSNLELKRKMQMQRDEEKNRRNNKDSTNQVMK
dr Tmc2b-2 VVLLMV-LAIYYLNAVSKAYQNSNLELKRKMQMQRDEEKNRRNNKDSTNQVMK
α10

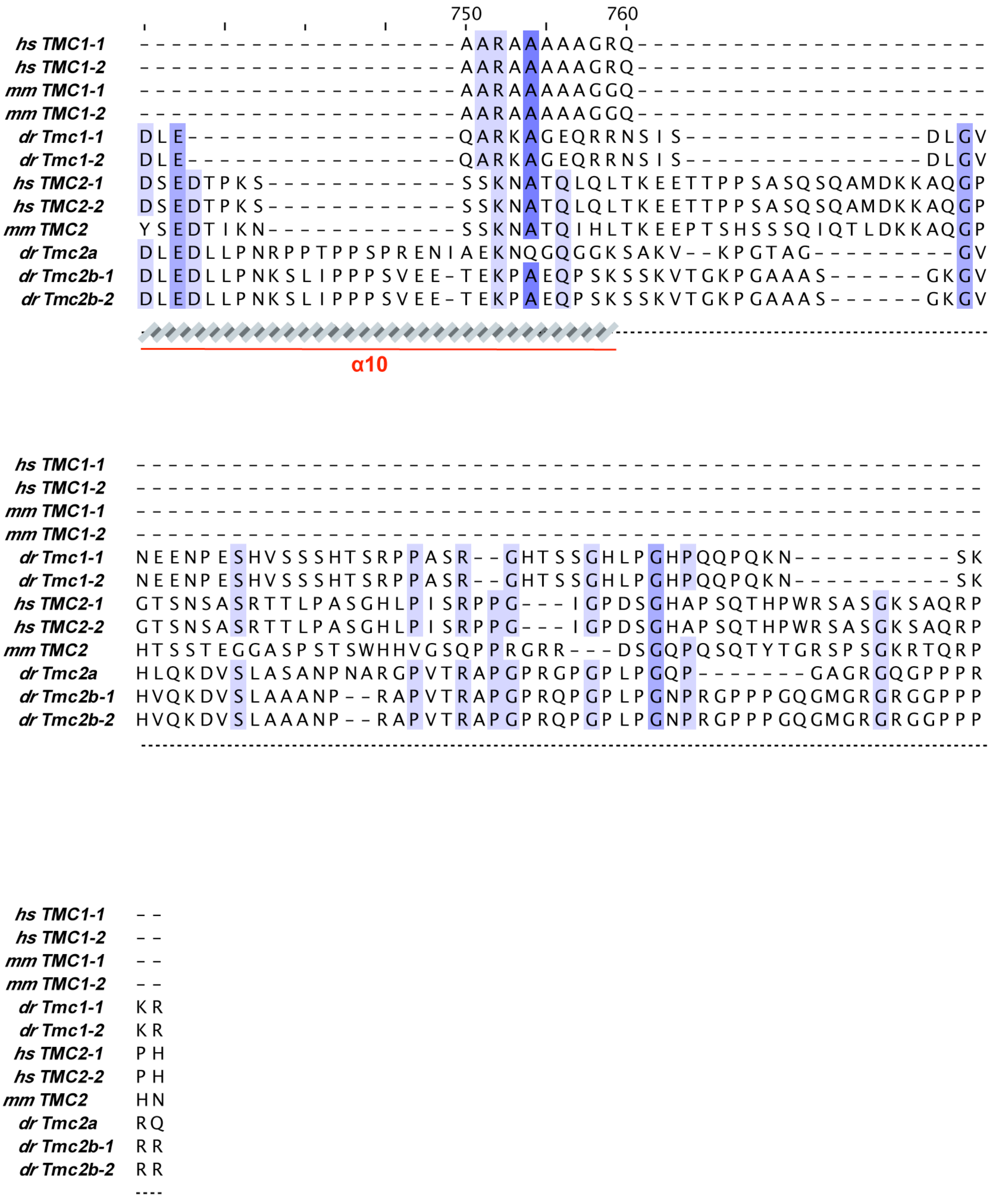


**Figure 7–figure Supplement 2**. **TMC sequence alignment.** Protein sequences correspond to: *hs* TMC1-1 (NCBI ID: NP_619636.2), *hs* TMC1-2 (NCBI ID: XP_016869745.1), *mm* TMC1-1 (NCBI ID: NP_083229.1), *mm* TMC1-2 (NCBI ID: XP_036017315.1), *dr* Tmc1-1 (NCBI ID: NP_001299610.1), *dr* Tmc1-2 (NCBI ID: XP_021331962.1), *hs* TMC2-1 (NCBI ID: NP_542789.2), *hs* TMC2-2 (NCBI ID: XP_005260717.1), *mm* TMC2 (NCBI ID: NP_619596.1), *dr* Tmc2a (NCBI ID: NP_001289166.1), *dr* Tmc2b-1 (NCBI ID: NP_001289152.1), and *dr* Tmc2b-2 (NCBI ID: XP_017211636.1). Secondary structures underneath the sequences are based on AF2 models of *hs* TMC1. Interface residues between TMC-NT/IL1 and CIB proteins were based on our AF2 model (for NT) and on Liang *et al*. (blue brackets).

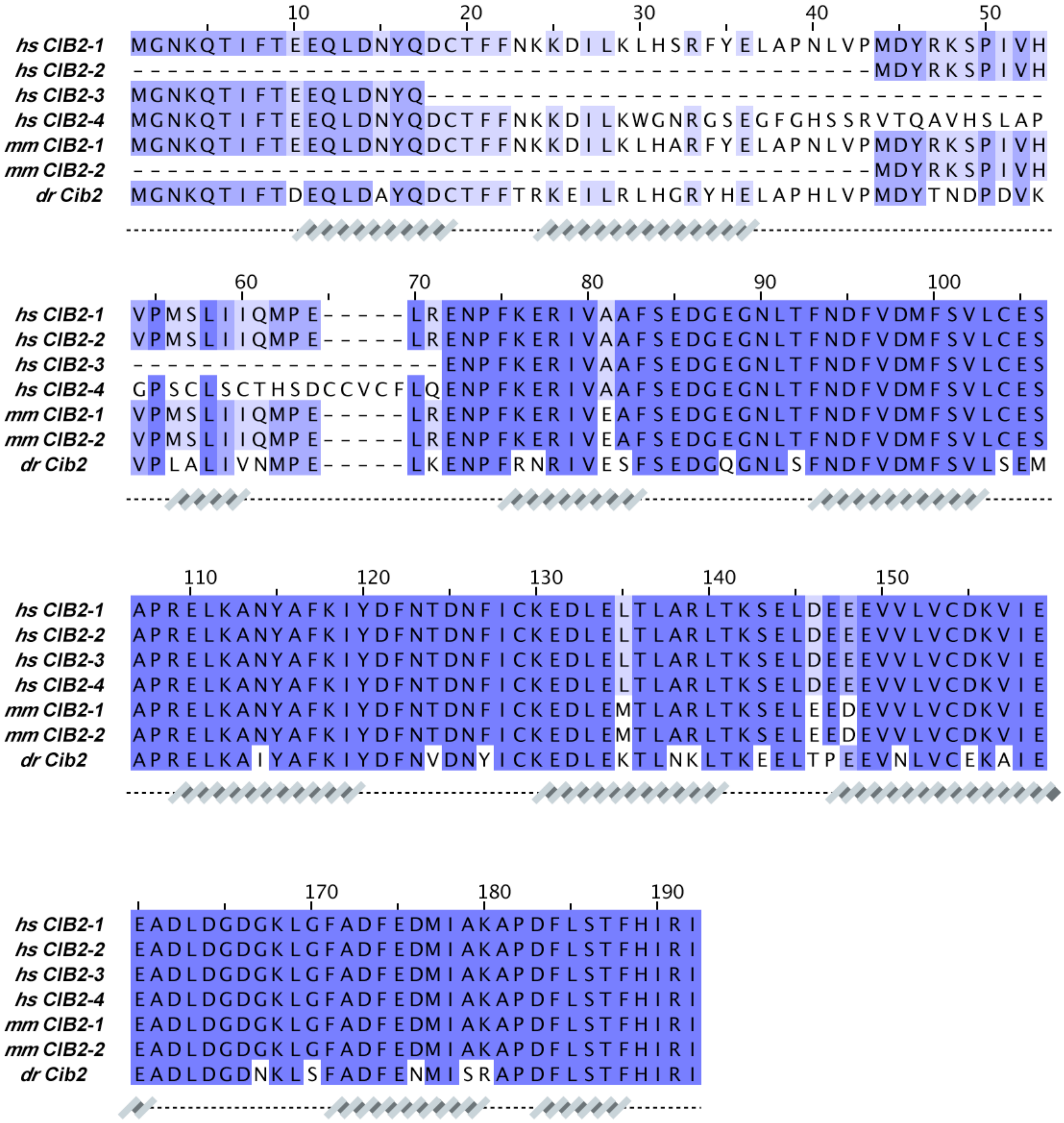


**Figure 7–figure supplement 3**. **CIB2 sequence alignment.** Protein sequences correspond to: *hs* CIB2-1 (NCBI ID: NP_006374.1) , *hs* CIB2-2 (NCBI ID: NP_001258817.1), *hs* CIB2-3 (NCBI ID: NP_001258818.1), *hs* CIB2-4 (NCBI ID: NP_001288153.1), *mm* CIB2-1 (NCBI ID: NP_062660.1), *mm* CIB2-2 (NCBI ID: XP_036011040.1), and *dr* Cib2 (NCBI ID: NP_957000.1). Secondary structures underneath the sequences are based on an AF2 model of *hs* CIB2.

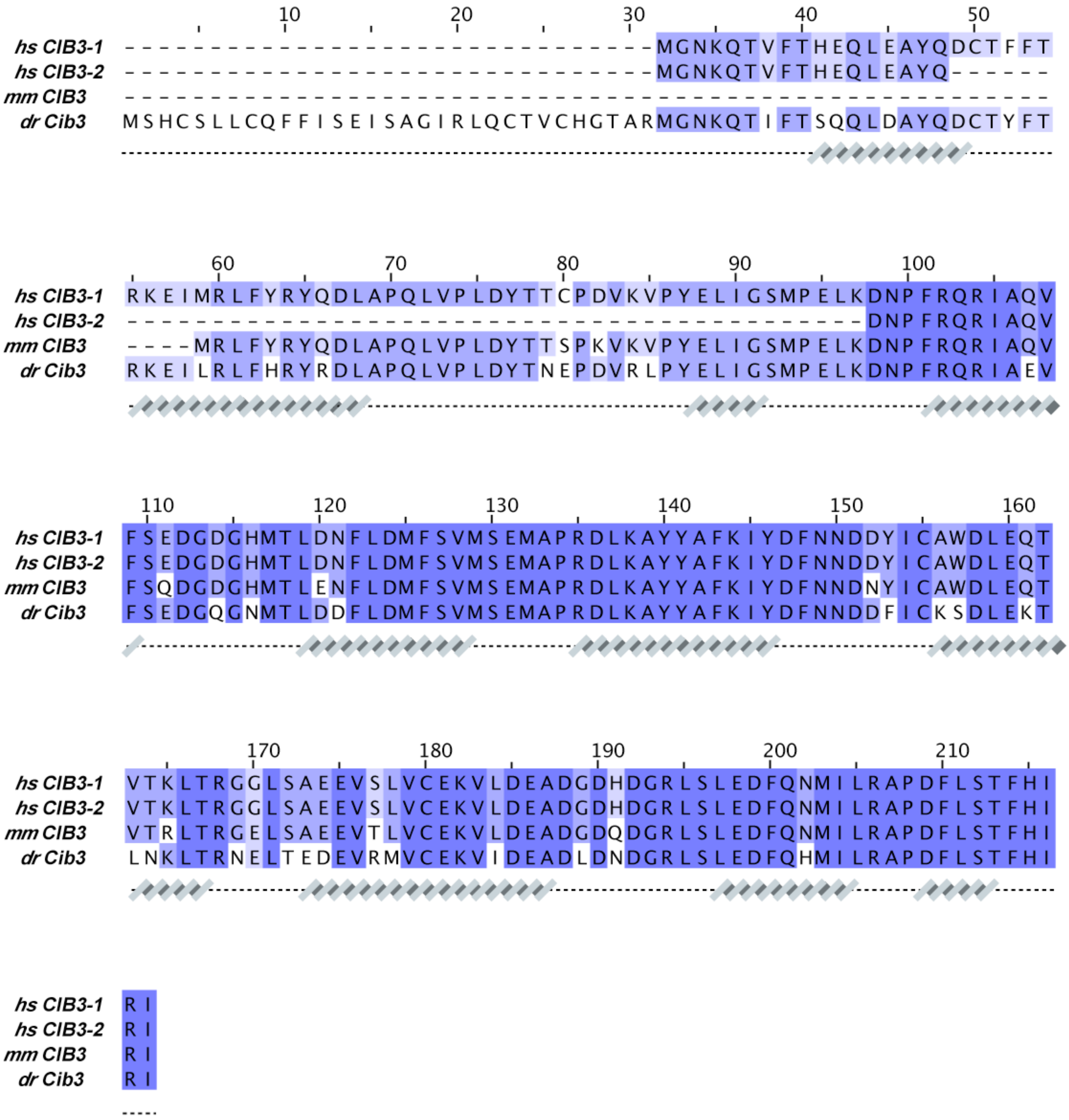


**Figure 7–figure supplement 4**. **CIB3 sequence alignment**. Protein sequences correspond to: *hs* CIB3-1 (NCBI ID: NP_473454.1), *hs* CIB3-2 (NCBI ID: NP_001287851.1), *mm* CIB3 (NCBI ID: NP_001074281.1), and *dr* Cib3 (NCBI ID: XP_003198013.2). Secondary structures underneath the sequences are based on an AF2 model of *hs* CIB3.

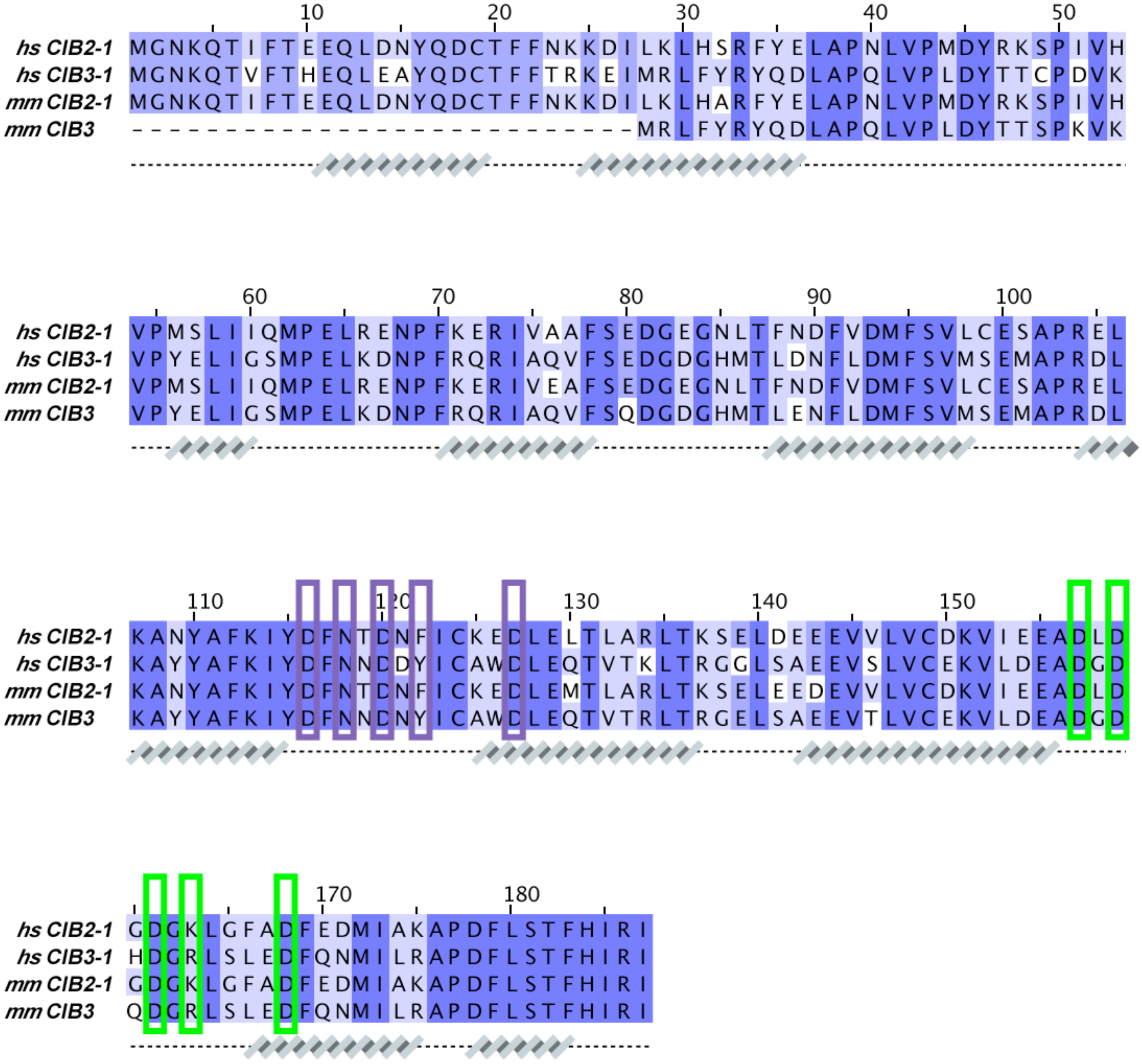

**Figure 7–figure supplement 5**. **CIB sequence alignment**. Protein sequences correspond to: *hs* CIB2-1 (NCBI ID: NP_006374.1), *hs* CIB3-1 (NCBI ID: NP_473454.1), *mm* CIB2-1 (NCBI ID: NP_062660.1), and *mm* CIB3 (NCBI ID: NP_001074281.1). Secondary structures underneath the sequences are based on AF2 models of *hs* CIB2/3. Cation-coordinating CIB residues were based on the X-ray crystal structure of *hs* CIB3 in complex with *mm* TMC1-IL1 bound to 2 $Mg^{2+}$ (PDB code: 6WUD; purple brackets for $Mg^{2+}$ bound to EF3 and green brackets for $Mg^{2+}$ bound to EF4).

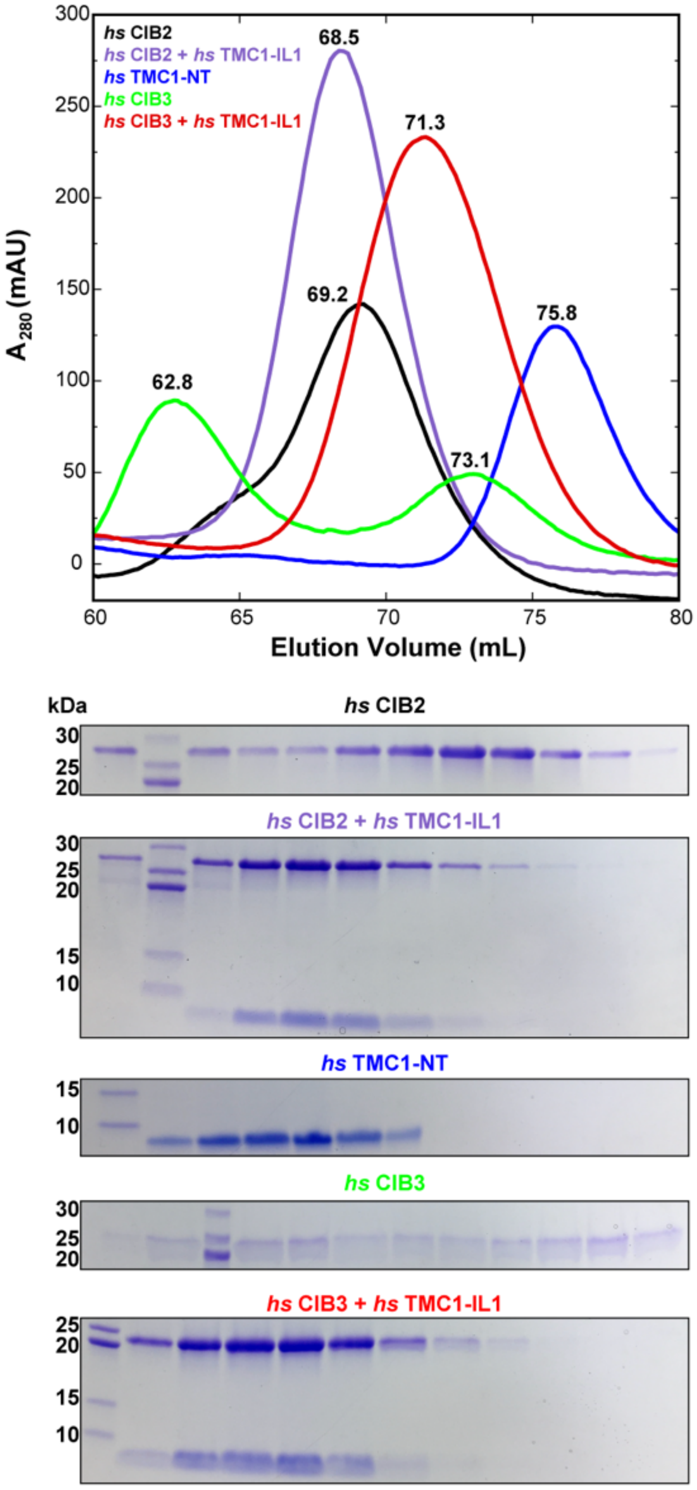


**Figure 7–figure supplement 6**. **SEC of *hs* CIB2 and *hs* CIB3 either refolded alone or co-refolded with *hs* TMC1-IL1, and *hs* TMC1-NT**. Representative traces for *hs* CIB2 (black), *hs* CIB3 (green), *hs* CIB2 + *hs* TMC1-IL1 (violet), *hs* CIB3 + *hs* TMC1-IL1 (red) and *hs* TMC1-NT (blue). Traces for *hs* CIB2, *hs* CIB2 + *hs* TMC1-IL1, *hs* CIB3 + *hs* TMC1-IL1, and *hs* TMC1-NT show peaks at 69.2 mL, 68.5 mL, 71.3 mL, and 75.8 mL, respectively. Trace for *hs* CIB3 showed two peaks at 62.8 mL and 73.1 mL. Experiments were performed using a Superdex S75 16/600 column in the presence of 3 mM $CaCl_2$. Coomassie-stained SDS-PAGE analyses of representative eluted fractions are shown below the chromatogram.

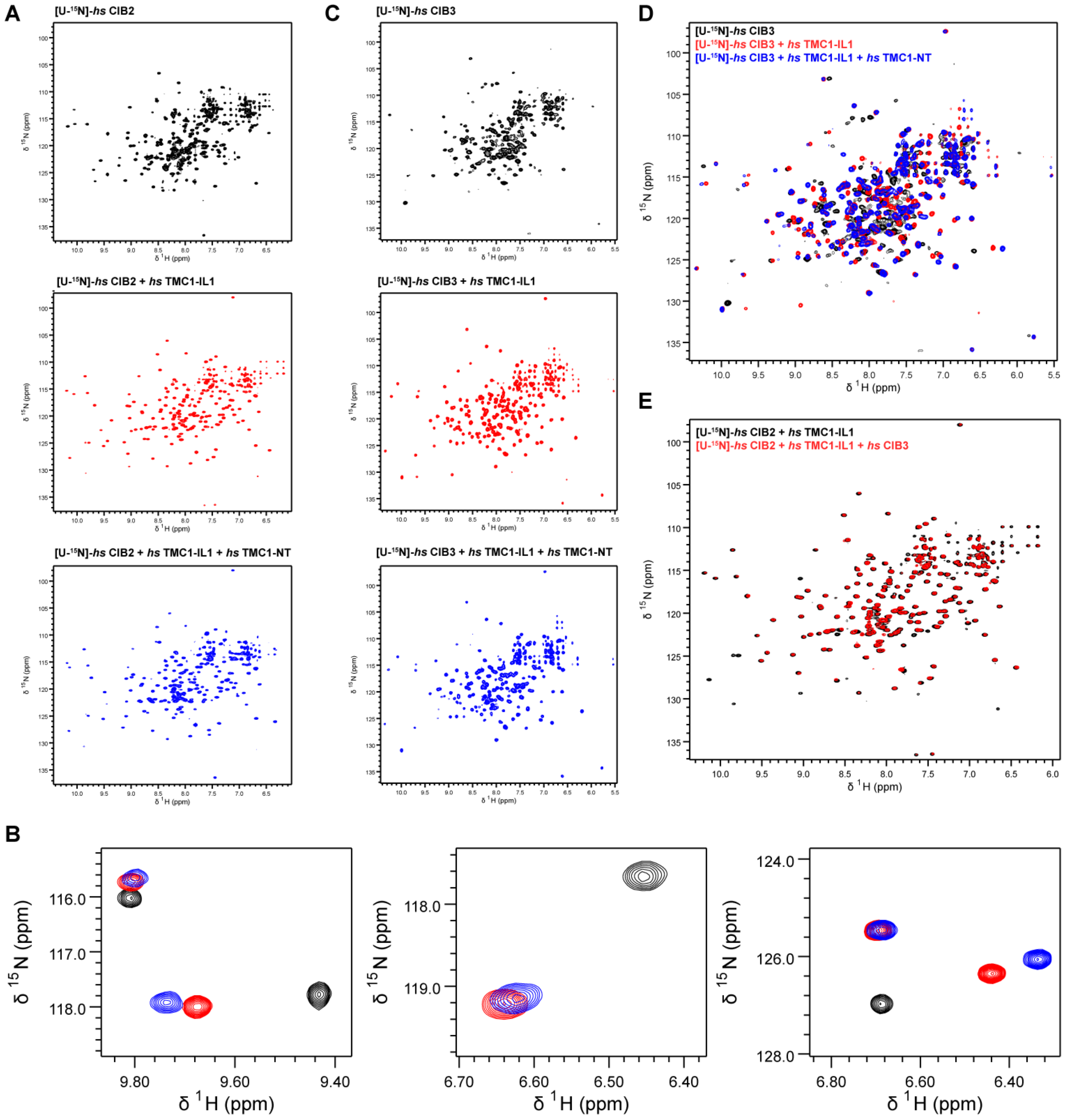


**Figure 7–figure supplement 7.** $^{1}$**H-**$^{15}$**N TROSY-HSQC spectra** of (**A**) *hs* CIB2, *hs* CIB2 + *hs* TMC1-IL1, and *hs* CIB2 + *hs* TMC1-IL1 + *hs* TMC1-NT individually (overlayed in Figure 7C). (**B**) Zoomed in spectra highlighting peak shifts that correspond to *hs* TMC1-IL1 and subsequent *hs* TMC1-NT-binding. (**C**,**D**) Spectra of *hs* CIB3, *hs* CIB3 + *hs* TMC1-IL1, *hs* CIB3 + *hs* TMC1-IL1 + *hs* TMC1-NT individually and overlayed, respectively. (**E**) Overlayed spectra of *hs* CIB2 + *hs* TMC1-IL1 and *hs* CIB2 + *hs* TMC1-IL1 + *hs* CIB3. NMR data were obtained in the presence of 3 mM $CaCl_2$. Some of these data are also shown in Figure 7.

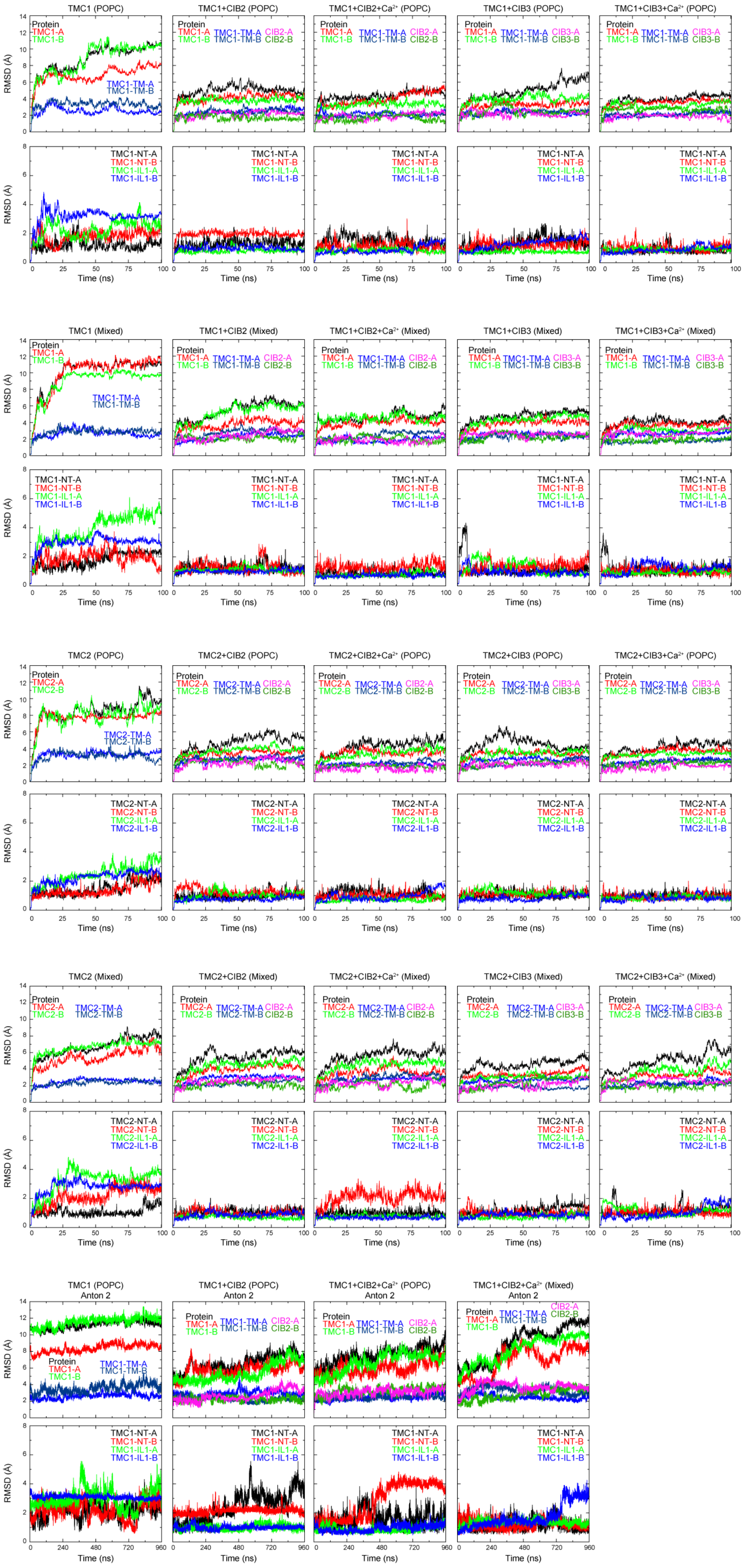
TMC1 (POPC)
TMC1+CIB2 (POPC)
TMC1+CIB2+Ca$^{2+}$ (POPC)
TMC1+CIB3 (POPC)
TMC1+CIB3+Ca$^{2+}$ (POPC)
TMC1 (Mixed)
TMC1+CIB2 (Mixed)
TMC1+CIB2+Ca$^{2+}$ (Mixed)
TMC1+CIB3 (Mixed)
TMC1+CIB3+Ca$^{2+}$ (Mixed)
TMC2 (POPC)
TMC2+CIB2 (POPC)
TMC2+CIB2+Ca$^{2+}$ (POPC)
TMC2+CIB3 (POPC)
TMC2+CIB3+Ca$^{2+}$ (POPC)
TMC2 (Mixed)
TMC2+CIB2 (Mixed)
TMC2+CIB2+Ca$^{2+}$ (Mixed)
TMC2+CIB3 (Mixed)
TMC2+CIB3+Ca$^{2+}$ (Mixed)
TMC1 (POPC) Anton 2
TMC1+CIB2 (POPC) Anton 2
TMC1+CIB2+Ca$^{2+}$ (POPC) Anton 2
TMC1+CIB2+Ca$^{2+}$ (Mixed) Anton 2
Protein
TMC1-A
TMC1-B
TMC1-TM-A
TMC1-TM-B
CIB2-A
CIB2-B
CIB3-A
CIB3-B
TMC1-NT-A
TMC1-NT-B
TMC1-IL1-A
TMC1-IL1-B
TMC2-A
TMC2-B
TMC2-TM-A
TMC2-TM-B
TMC2-NT-A
TMC2-NT-B
TMC2-IL1-A
TMC2-IL1-B
RMSD (Å)
Time (ns)

**Figure 8–figure supplement 1**. **Stability of TMC + CIB complexes, monomers, and subdomains**. RMSDs of the full protein (ALL), TMC monomers A and B (TMC1/2-A and TMC1/2-B), transmembrane helices of TMC monomers A and B (TMC1/2-TM-A and TMC1/2-TM-B), and CIB monomers are shown on top for each simulated system as labeled. RMSDs for TMC-NT and -IL1 fragments are shown on bottom plots. RMSD values were calculated for Cα atoms after alignment to the initial frame of each trajectory. For Anton 2 trajectories, RMSD values were calculated for Cα atoms after alignment to the initial frame of their respective NAMD trajectory.

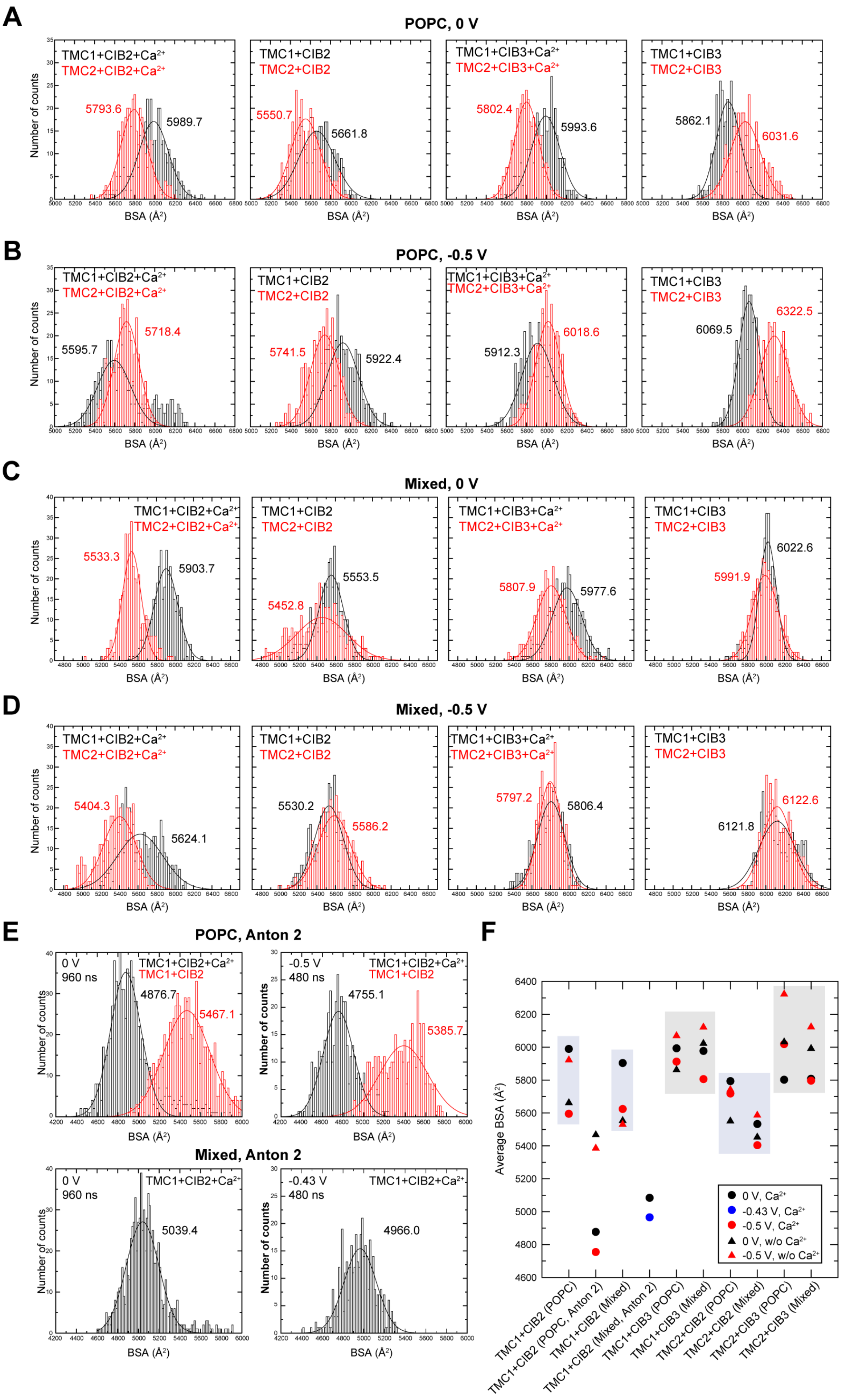
A
POPC, 0 V
TMC1+CIB2+Ca²⁺
TMC2+CIB2+Ca²⁺
5793.6
5989.7
TMC1+CIB2
TMC2+CIB2
5550.7
5661.8
TMC1+CIB3+Ca²⁺
TMC2+CIB3+Ca²⁺
5802.4
5993.6
TMC1+CIB3
TMC2+CIB3
5862.1
6031.6
Number of counts
BSA (Å²)
B
POPC, -0.5 V
5595.7
5718.4
5741.5
5922.4
5912.3
6018.6
6069.5
6322.5
C
Mixed, 0 V
5533.3
5903.7
5452.8
5553.5
5807.9
5977.6
5991.9
6022.6
D
Mixed, -0.5 V
5404.3
5624.1
5530.2
5586.2
5797.2
5806.4
6121.8
6122.6
E
POPC, Anton 2
0 V
960 ns
TMC1+CIB2+Ca²⁺
TMC1+CIB2
4876.7
5467.1
-0.5 V
480 ns
4755.1
5385.7
Mixed, Anton 2
5039.4
-0.43 V
4966.0
F
Average BSA (Å²)
0 V, Ca²⁺
-0.43 V, Ca²⁺
-0.5 V, Ca²⁺
0 V, w/o Ca²⁺
-0.5 V, w/o Ca²⁺
TMC1+CIB2 (POPC)
TMC1+CIB2 (POPC, Anton 2)
TMC1+CIB2 (Mixed)
TMC1+CIB2 (Mixed, Anton 2)
TMC1+CIB3 (POPC)
TMC1+CIB3 (Mixed)
TMC2+CIB2 (POPC)
TMC2+CIB2 (Mixed)
TMC2+CIB3 (POPC)
TMC2+CIB3 (Mixed)

**Figure 8–figure supplement 2**. **BSA in *hs* TMC1/2 and *hs* CIB2/3 complexes**. (**A**) Histogram plots for BSA values computed for POPC systems during equilibrium simulations (10-100 ns). (**B**) Histogram plots for BSA values computed for POPC systems during simulations at -0.5 V (100-200 ns). (**C**) Histogram plots for BSA values computed for mixed membrane systems during equilibrium simulations (10-100 ns). (**D**) Histogram plots for BSA values computed for mixed membrane systems during simulations at -0.5 V (100-200 ns). (**E**) Histogram plots for BSA values computed for POPC and mixed membrane systems during Anton 2 simulations at 0 V (0-960 ns), -0.43 V (240-720 ns for mixed system), and -0.5 V (and 960-1440 ns for POPC systems). BSA count distributions for all plots were fitted to a normal distribution with mean BSA values labeled with corresponding colors. (**F**) Summary of mean BSA values.

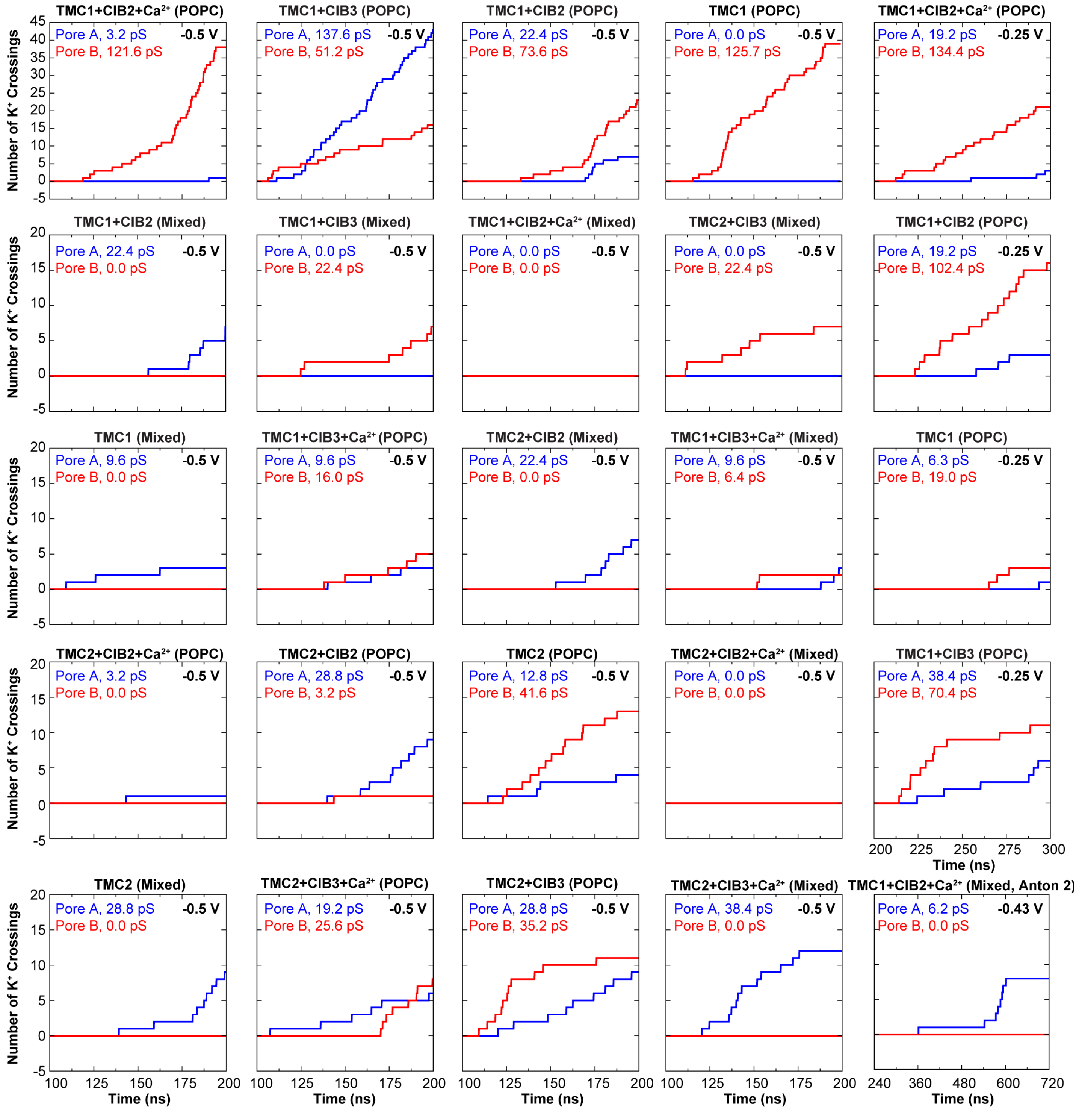


**Figure 8–figure supplement 3**. **Number of $K^+$ crossings as a function of time for pores A and B of *hs* TMC1 and *hs* TMC2**. Plots for systems simulated at -0.5 V are shown to start after 100 ns of equilibration and lasting 100 ns of simulation (200 ns total). Plots for systems simulated at -0.25 V are shown to start after 100 ns of equilibration + 100 ns of simulation at -0.5 V (200 ns) and lasting 100 ns (300 ns total). Plot for system simulated in Anton 2 shown to start after 240 ns of equilibration + 480 ns of simulation at -0.43 V (720 ns total). Conductance values calculated for $K^+$ crossings.

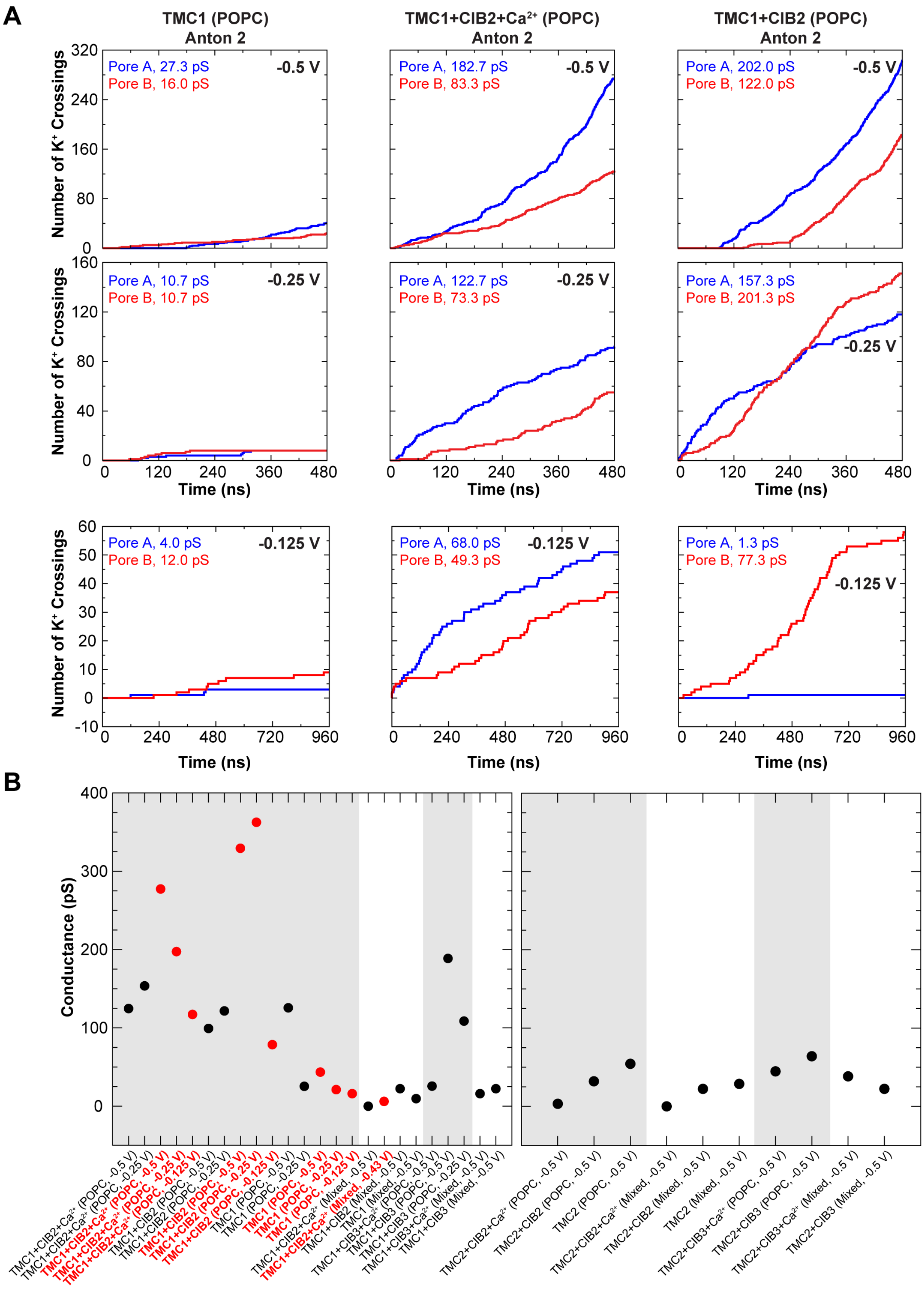


**Figure 8–figure supplement 4. Conduction events in long timescale simulations and summary of predicted conductance values.** (**A**) Number of $K^+$ crossings as a function of time for pores A and B of *hs* TMC1 simulated in Anton 2. Plots are concatenated in Figure 7H. (**B**) Summary of predicted conductance values obtained from all simulations of

TMC1 (left) and TMC2 (right). Gray shadow denotes POPC systems. Red circles are for Anton 2 simulations. Conductance values for each system were plotted as a sum of both pores. Conductance values calculated for $K^+$ crossings.

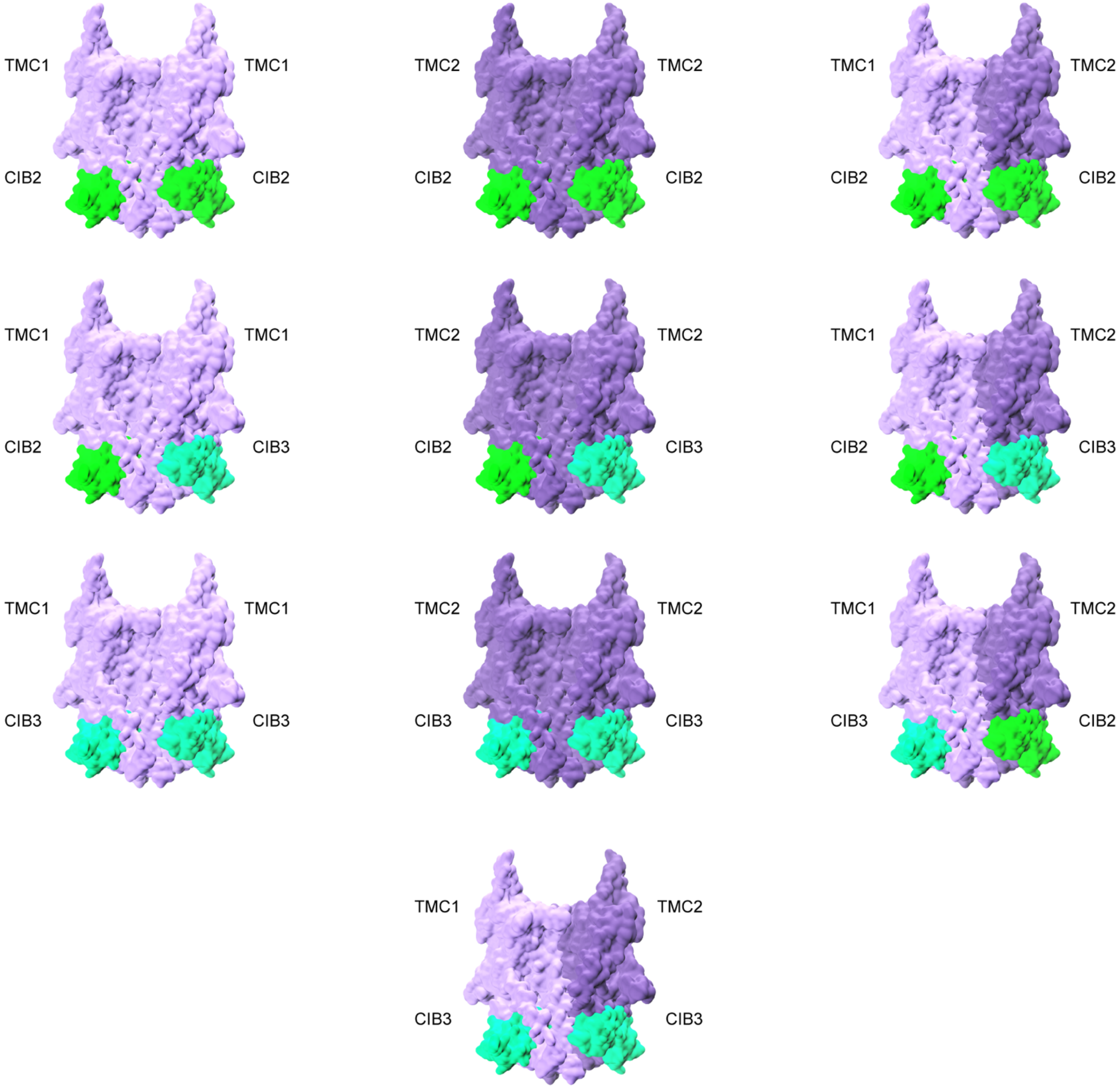


**Figure 8–figure supplement 5**. **Possible combinations of TMC1, TMC2, CIB2, and CIB3 complexes**.

**Figure 8—table supplement 1. Summary of simulations**

| System | Label | $Ca^{2+}$ | Membrane | Start | Length (ns) | Voltage (V) | Size (# of atoms) | Initial Size ($nm^3$) |
|---|---|---|---|---|---|---|---|---|
| *hs* TMC1 + *hs* CIB2 | S1a | Yes | POPC | - | 100 | 0† | 395,283 | 16.1 x 16.1 x 16.6 |
| | S1b | | | S1a | 100 | -0.5 | | |
| | S1c | | | S1b | 100 | -0.25 | | |
| | S1d | | | S1a | 960 | 0 | | |
| | S1e | | | S1d | 480 | -0.5 | | |
| | S1f | | | S1e | 480 | -0.25 | | |
| | S1g | | | S1f | 960 | -0.125 | | |
| *hs* TMC1 + *hs* CIB2 | S2a | No | POPC | - | 100 | 0† | 395,200 | 16.1 x 16.1 x 16.6 |
| | S2b | | | S2a | 100 | -0.5 | | |
| | S2c | | | S2b | 100 | -0.25 | | |
| | S2d | | | S2a | 960 | 0 | | |
| | S2e | | | S2d | 480 | -0.5 | | |
| | S2f | | | S2e | 480 | -0.25 | | |
| | S2g | | | S2f | 960 | -0.125 | | |
| *hs* TMC1 | S3a | - | POPC | - | 100 | 0† | 395,146 | 16.1 x 16.1 x 16.6 |
| | S3b | | | S3a | 99.25 | -0.5 | | |
| | S3c | | | S3b | 100.875 | -0.25 | | |
| | S3d | | | S3a | 960 | 0 | | |
| | S3e | | | S3d | 480 | -0.5 | | |
| | S3f | | | S3e | 480 | -0.25 | | |
| | S3g | | | S3f | 960 | -0.125 | | |
| *hs* TMC1 + *hs* CIB2 | S4a | Yes | Mixed[a] | - | 100 | 0† | 425,170 | 16.8 x 16.7 x 16.6 |
| | S4b | | | S4a | 100 | -0.5 | | |
| | S4c | | | S4a | 240 | 0 | | |
| | S4d | | | S4c | 480 | -0.43 | | |
| | S5e | | | S4c | 720 | 0 | | |
| *hs* TMC1 + *hs* CIB2 | S5a | No | Mixed[a] | - | 100 | 0† | 424,916 | 16.8 x 16.7 x 16.6 |
| | S5b | | | S5a | 100 | -0.5 | | |
| *hs* TMC1 | S6a | - | Mixed[a] | - | 100 | 0† | 425,224 | 16.8 x 16.8 x 16.6 |
| | S6b | | | S6a | 100 | -0.5 | | |
| *hs* TMC1 + *hs* CIB3 | S7a | Yes | POPC | - | 100 | 0† | 394,843 | 16.1 x 16.1 x 16.6 |
| | S7b | | | S7a | 100 | -0.5 | | |
| *hs* TMC1 + *hs* CIB3 | S8a | No | POPC | - | 100 | 0† | 394,750 | 16.1 x 16.1 x 16.6 |
| | S8b | | | S8a | 100 | -0.5 | | |
| | S8c | | | S8b | 100 | -0.25 | | |
| *hs* TMC1 + *hs* CIB3 | S9a | Yes | Mixed[a] | - | 100 | 0† | 424,929 | 16.8 x 16.8 x 16.6 |
| | S9b | | | S9a | 100 | -0.5 | | |
| *hs* TMC1 + *hs* CIB3 | S10a | No | Mixed[a] | - | 100 | 0† | 425,005 | 16.8 x 16.7 x 16.6 |
| | S10b | | | S10a | 100 | -0.5 | | |
| *hs* TMC2 + *hs* CIB2 | S11a | Yes | POPC | - | 100 | 0† | 434,265 | 16.8 x 16.9 x 16.7 |
| | S11b | | | S11a | 100 | -0.5 | | |
| *hs* TMC2 + *hs* CIB2 | S12a | No | POPC | - | 100 | 0† | 433,630 | 16.9 x 16.9 x 16.6 |
| | S12b | | | S12a | 100 | -0.5 | | |
| *hs* TMC2 | S13a | - | POPC | - | 100 | 0† | 433,327 | 16.9 x 16.9 x 16.7 |
| | S13b | | | S13a | 100 | -0.5 | | |
| *hs* TMC2 + *hs* CIB2 | S14a | Yes | Mixed[a] | - | 100 | 0† | 427,357 | 16.8 x 16.8 x 16.5 |
| | S14b | | | S14a | 100 | -0.5 | | |
| *hs* TMC2 + *hs* CIB2 | S15a | No | Mixed[a] | - | 100 | 0† | 427,254 | 16.8 x 16.8 x 16.5 |
| | S15b | | | S15a | 100 | -0.5 | | |
| *hs* TMC2 | S16a | - | Mixed[a] | - | 100 | 0† | 428,031 | 16.8 x 16.8 x 16.6 |
| | S16b | | | S16a | 100 | -0.5 | | |
| *hs* TMC2 + *hs* CIB3 | S17a | Yes | POPC | - | 100 | 0† | 429,479 | 16.8 x 16.8 x 16.6 |
| | S17b | | | S17a | 100 | -0.5 | | |
| *hs* TMC2 + *hs* CIB3 | S18a | No | POPC | - | 100 | 0† | 429,600 | 16.9 x 16.8 x 16.6 |
| | S18b | | | S18a | 100 | -0.5 | | |
| *hs* TMC2 + *hs* CIB3 | S19a | Yes | Mixed[a] | - | 100 | 0† | 428,972 | 16.8 x 16.8 x 16.6 |
| | S19b | | | S19a | 100 | -0.5 | | |
| *hs* TMC2 + *hs* CIB3 | S20a | No | Mixed[a] | - | 100 | 0† | 428,904 | 16.8 x 16.9 x 16.6 |
| | S20b | | | S20a | 100 | -0.5 | | |
| | | | | Total time | 14,480.125 | | | |

*a* lipid membrane bilayer was comprised of 65% POPC, 30% cholesterol and 5% PIP2. † denotes equilibrium simulations that consisted of 2,000 steps of minimization, 0.5 ns of dynamics with everything excluding lipid tails fixed/constrained, 0.5 ns of dynamics with harmonic constraints applied to the protein (1 kcal $mol^{-1}$ $Å^{-2}$), 1 ns of free dynamics in the *NpT* ensemble ($\gamma$ = 1 $ps^{-1}$), and up to 100 ns of free dynamics in the *NpT* ensemble ($\gamma$ = 0.1 $ps^{-1}$).

| Figure 8—table supplement 2. Ion conduction | | | | | | | | | |
|---|---|---|---|---|---|---|---|---|---|
| Label | Length (ns) | Voltage (V) | $K^+$ (A) | $K^+$ (B) | $Cl^-$ (A) | $Cl^-$ (B) | I† (pA) | C† (pS) | BSA (Å$^2$) |
| S1a | 100 | 0 | - | - | - | - | - | - | 5,989.7 |
| S1b | 100 | -0.5 | 1 | 38 | 0 | 0 | 62.4 | 124.8 | 5,595.7 |
| S1c | 100 | -0.25 | 3 | 21 | 0 | 0 | 38.4 | 153.6 | - |
| S1d | 960 | 0 | - | - | - | - | - | - | 4,876.7 |
| S1e | 480 | -0.5 | 274 | 125 | 17 | 0 | 138.7 | 277.3 | 4,755.1 |
| S1f | 480 | -0.25 | 92 | 55 | 1 | 0 | 49.3 | 197.3 | - |
| S1g | 960 | -0.125 | 51 | 37 | 0 | 0 | 14.7 | 117.3 | - |
| S2a | 100 | 0 | - | - | - | - | - | - | 5,661.8 |
| S2b | 100 | -0.5 | 7 | 23 | 0 | 1 | 49.6 | 99.2 | 5,922.4 |
| S2c | 100 | -0.25 | 3 | 16 | 0 | 0 | 30.4 | 121.6 | - |
| S2d | 960 | 0 | - | - | - | - | - | - | 5,467.1 |
| S2e | 480 | -0.5 | 303 | 183 | 6 | 2 | 164.7 | 329.3 | 5,385.7 |
| S2f | 480 | -0.25 | 118 | 151 | 1 | 1 | 90.3 | 361.3 | - |
| S2g | 960 | -0.125 | 1 | 58 | 0 | 0 | 9.8 | 78.7 | - |
| S3a | 100 | 0 | - | - | - | - | - | - | - |
| S3b | 99.25 | -0.50 | 0 | 39 | 0 | 0 | 62.9 | 125.7 | - |
| S3c | 100.875 | -0.25 | 1 | 3 | 0 | 0 | 6.3 | 25.4 | - |
| S3d | 960 | 0 | - | - | - | - | - | - | - |
| S3e | 480 | -0.5 | 41 | 24 | 0 | 0 | 21.7 | 43.4 | - |
| S3f | 480 | -0.25 | 8 | 8 | 0 | 0 | 5.3 | 21.3 | - |
| S3g | 960 | -0.125 | 3 | 9 | 0 | 0 | 2.0 | 16.0 | - |
| S4a | 100 | 0 | - | - | - | - | - | - | 5,903.7 |
| S4b | 100 | -0.50 | 0 | 0 | 0 | 0 | 0.0 | 0.0 | 5,624.1 |
| S4c | 240 | 0 | - | - | - | - | - | - | 5,084.2 |
| S4d | 480 | -0.43 | 8 | 0 | 0 | 0 | 2.7 | 6.2 | 4,966.0 |
| S4e | 720 | 0 | - | - | - | - | - | - | 5,039.4 |
| S5a | 100 | 0 | - | - | - | - | - | - | 5,553.5 |
| S5b | 100 | -0.50 | 7 | 0 | 0 | 0 | 11.2 | 22.4 | 5,530.2 |
| S6a | 100 | 0 | - | - | - | - | - | - | - |
| S6b | 100 | -0.50 | 3 | 0 | 0 | 0 | 4.8 | 9.6 | - |
| S7a | 100 | 0 | - | - | - | - | - | - | 5,993.6 |
| S7b | 100 | -0.50 | 3 | 5 | 0 | 0 | 12.8 | 25.6 | 5,912.3 |
| S8a | 100 | 0 | - | - | - | - | - | - | 5,862.1 |
| S8b | 100 | -0.50 | 43 | 16 | 0 | 0 | 94.4 | 188.8 | 6,069.5 |
| S8c | 100 | -0.25 | 6 | 11 | 0 | 0 | 27.2 | 108.8 | - |
| S9a | 100 | 0 | - | - | - | - | - | - | 5,977.6 |
| S9b | 100 | -0.50 | 3 | 2 | 0 | 0 | 8.0 | 16.0 | 5,806.4 |
| S10a | 100 | 0 | - | - | - | - | - | - | 6,022.6 |
| S10b | 100 | -0.50 | 0 | 7 | 0 | 0 | 11.2 | 22.4 | 6,121.8 |
| S11a | 100 | 0 | - | - | - | - | - | - | 5,793.6 |
| S11b | 100 | -0.50 | 1 | 0 | 0 | 0 | 1.6 | 3.2 | 5,718.4 |
| S12a | 100 | 0 | - | - | - | - | - | - | 5,550.7 |
| S12b | 100 | -0.50 | 9 | 1 | 0 | 0 | 16.0 | 32.0 | 5,741.5 |
| S13a | 100 | 0 | - | - | - | - | - | - | - |
| S13b | 100 | -0.50 | 4 | 13 | 0 | 0 | 27.2 | 54.4 | - |
| S14a | 100 | 0 | - | - | - | - | - | - | 5,533.3 |
| S14b | 100 | -0.50 | 0 | 0 | 0 | 0 | 0.0 | 0.0 | 5,404.3 |
| S15a | 100 | 0 | - | - | - | - | - | - | 5,452.8 |
| S15b | 100 | -0.50 | 7 | 0 | 0 | 0 | 11.2 | 22.4 | 5,586.2 |
| S16a | 100 | 0 | - | - | - | - | - | - | - |
| S16b | 100 | -0.50 | 9 | 0 | 0 | 0 | 14.4 | 28.8 | - |
| S17a | 100 | 0 | - | - | - | - | - | - | 5,802.4 |
| S17b | 100 | -0.50 | 6 | 8 | 0 | 0 | 22.4 | 44.8 | 6,018.6 |
| S18a | 100 | 0 | - | - | - | - | - | - | 6,031.6 |
| S18b | 100 | -0.50 | 9 | 11 | 0 | 0 | 32.0 | 64.0 | 6,322.5 |
| S19a | 100 | 0 | - | - | - | - | - | - | 5,807.9 |
| S19b | 100 | -0.50 | 12 | 0 | 0 | 0 | 19.2 | 38.4 | 5,797.2 |
| S20a | 100 | 0 | - | - | - | - | - | - | 5,991.9 |
| S20b | 100 | -0.50 | 0 | 7 | 0 | 0 | 11.2 | 22.4 | 6,122.6 |

† Currents, conductance, and average BSA values are reported for combined monomers A and B.
